# Discovery of the first selective M_4_ muscarinic acetylcholine receptor antagonists with *in vivo* anti-parkinsonian and anti-dystonic efficacy

**DOI:** 10.1101/2020.10.12.324152

**Authors:** Mark S. Moehle, Aaron M. Bender, Jonathan W. Dickerson, Daniel J. Foster, Yuping Donsante, Weimin Peng, Zoey Bryant, Thomas M. Bridges, Sichen Chang, Katherine J. Watson, Jordan C. O’Neill, Julie L. Engers, Li Peng, Alice L. Rodriguez, Colleen M. Niswender, Craig W. Lindsley, Ellen J. Hess, P. Jeffrey Conn, Jerri M. Rook

**Affiliations:** Department of Pharmacology, Warren Center for Neuroscience Drug Discovery, Vanderbilt University, Nashville, TN, 37232, United States; Department of Pharmacology, Warren Center for Neuroscience Drug Discovery, and Vanderbilt Kennedy Center, Vanderbilt University, Nashville, TN, 37232, United States; Department of Pharmacology & Chemical Biology, Emory University, Atlanta, Georgia, 30322, United States; Department of Pharmacology, Warren Center for Neuroscience Drug Discovery, Vanderbilt University, Nashville, TN 37232, United States; Departments of Pharmacology and Chemistry, Warren Center for Neuroscience Drug Discovery, Vanderbilt University, Nashville, TN, 37232, United States

**Keywords:** acetylcholine, muscarinic, Parkinson disease, dopamine, cholinergic, dystonia

## Abstract

Non-selective antagonists of muscarinic acetylcholine receptors (mAChRs) that broadly inhibit all five mAChR subtypes provide an efficacious treatment for some movement disorders, including Parkinson disease and dystonia. Despite their efficacy in these and other central nervous system disorders, anti-muscarinic therapy has limited utility due to severe adverse effects that often limit their tolerability by patients. Recent advances in understanding the roles that each mAChR subtype plays in disease pathology suggest that highly selective ligands for individual subtypes may underlie the anti-parkinsonian and anti-dystonic efficacy observed with the use of non-selective anti-muscarinic therapeutics. Our recent work has indicated that the M_4_ muscarinic acetylcholine receptor has several important roles in opposing aberrant neurotransmitter release, intracellular signaling pathways, and brain circuits associated with movement disorders. This raises the possibility that selective antagonists of M_4_ may recapitulate the efficacy of non-selective anti-muscarinic therapeutics and may decrease or eliminate the adverse effects associated with these drugs. However, this has not been directly tested due to lack of selective antagonists of M_4_. Here we utilize genetic mAChR knockout animals in combination with non-selective mAChR antagonists to confirm that the M_4_ receptor underlies the locomotor-stimulating and anti-parkinsonian efficacy in rodent models. We also report the synthesis, discovery, and characterization of the first-in-class selective M_4_ antagonists VU6013720, VU6021302, and VU6021625 and confirm that these optimized compounds have anti-parkinsonian and anti-dystonic efficacy in pharmacological and genetic models of movement disorders.

## Introduction

Dopamine (DA) release and signaling in the basal ganglia (BG) are critical for fine-tuned motor control and locomotor ability^1–4^. When DA release or signaling is diminished, such as in Parkinson disease (PD) following the death of DA releasing cells or in genetic forms of dystonia, aberrant motor behaviors are present^1–6^. In many of these disease states, especially PD, treatment often centers around boosting DA levels in the brain through administration of the dopamine prodrug levodopa (L-DOPA), preventing the breakdown of DA, or directly activating DA receptors^7^. However, these treatments can often lead to severe side effects such as dyskinesia, are not effective at treating all the symptoms in the disease state, and their efficacy is unreliable with disease progression and after chronic use^8^. Development of non-DA based therapies with different mechanisms of action that do not directly target the DA system could meet a large unmet clinical need in several movement disorders^9,10^.

One possible non-DA based treatment mechanism is through targeting of muscarinic acetylcholine receptors (mAChRs)^11,12^. Acetylcholine (ACh) acting through mAChRs has powerful neuromodulatory actions on the BG motor circuit^10,13,14^. Activation of mAChRs induces several actions to oppose DA release, DA signaling, as well as related motor behaviors. Consistent with these multiple actions on DA release and signaling, anti-mAChR therapy that targets each of the five mAChR subtypes (M_1_-M_5_) equally has efficacy in reducing the primary motor symptoms of PD and dystonia^7,9,11–13,15–19^. However, like DA-targeted therapies, despite their efficacy, non-selective anti-mAChR therapy can lead to serious on-target adverse effects that limits their tolerability by patients^11,20^.

Recent pharmacological and genetic studies have made it possible to define unique roles of individual mAChR subtypes in motor disorders^10,17^. These data raise the possibility that targeting of individual mAChR subtypes with truly selective and specific pharmacological ligands may maintain the efficacy but eliminate or reduce the adverse effects associated with non-selective mAChR pharmacological agents^10^. Several of the peripheral adverse effects associated with non-selective anti-muscarinic side effects are likely mediated by M_2_ and M_3_, and central side effects centering around memory and cognition may be due to M_1_^9,10,17,21,22^. However, recent studies using first-in-class M_4_ positive allosteric modulators (PAMs) along with genetic approaches, suggest that potentiation of M_4_ signaling opposes DA signaling in the BG motor circuit via multiple mechanisms. For example, potentiation of M_4_ activation in the dorsal striatum can cause a sustained inhibition of DA release^19^. Furthermore, M_4_ potentiation on BG direct pathway terminals in the substantia nigra pars reticulata (SNr), the primary BG output nucleus in rodents, directly opposes dopamine receptor subtype 1 (D_1_) signaling in these cells, leading to tonic inhibition of BG direct pathway activity and reduced locomotion^15^. Genetic deletion of M_4_ either globally or in D_1_ DA expressing spiny projection neurons (D_1_-SPNs), which form the BG direct pathway, recapitulates many of our pharmacological findings as well^16,18^. Mice with selective M_4_ deletion in D_1_-SPNs or globally are also more sensitive to the pro-locomotor effects of psychomotor stimulants, have elevated basal DA levels, and are hyperlocomotive^16,18^. These genetic and pharmacological studies of mAChR subtypes suggest that M_4_ may be the dominant mAChR in the regulation of DA and locomotor activity, and that M_4_ selective inhibitors may maintain the efficacy seen with non-selective anti-mAChR therapeutics in movement disorders while reducing or eliminating their side effects^9^.

Directly testing the hypothesis that M_4_ antagonism will have efficacy in movement disorders has been greatly limited due to the lack of truly selective M_4_ inhibitors^9^. To evaluate whether M_4_ underlies the efficacy of non-selective mAChR antagonists, we report the discovery of a first-in-class series of M_4_ selective antagonists. Utilizing knockout (KO) animals in conjunction with these novel, selective M_4_ antagonists, our data support the role of M_4_ in movement disorders and demonstrate anti-parkinsonian and anti-dystonic efficacy of M_4_ antagonists in preclinical models.

## Results

### M_4_ underlies the anti-parkinsonian and pro-locomotor efficacy of anti-mAChR compounds

To determine which mAChR subtype underlies the locomotor-stimulating effects of the anti-mAChR agent scopolamine, we first assessed the effects of multiple doses of scopolamine on locomotor activity in wildtype (WT) mice^23^. Mice were placed into an open field chamber and allowed to habituate for 90 minutes while activity (total distance traveled in cm/5 minute bins) was recorded. After 90 minutes, scopolamine (0.1 – 3 mg/kg, 10 ml/kg 10% Tween 80, intraperitoneal (i.p.)) was injected and total distance traveled was recorded (Figure 1A). Scopolamine increased total distance traveled from 90 – 150 minutes in a dose dependent manner, with the 3 mg/kg dose showing maximal efficacy (Figure 1A, B; One-way ANOVA with Dunnett’s post-hoc test; F_4,25_=9.064; p < 0.0001). The administration of 0.1 or 0.3 mg/kg of scopolamine did not significantly increase locomotor activity (1346.0 + 385.9 cm for vehicle, 509.7 + 193.5 cm for 0.1 mg/kg scopolamine, 1322.0 + 324.9 cm for 0.3 mg/kg scopolamine; Figure 1A, B; p > 0.05). In contrast, 1 and 3 mg/kg of scopolamine induced a significant increase in locomotor activity from 90 – 150 minutes (1346.0 + 385.9 cm for vehicle, 5272.0 + 726.4 cm for 1 mg/kg scopolamine, 6796.0 + 1870.0 cm for 3 mg/kg scopolamine; Figure 1A, B; p < 0.05 for 1 mg/kg; p < 0.01 for 3 mg/kg). We then repeated this procedure with 3 mg/kg scopolamine in M_1_ or M_4_ global KO mice and compared responses to their littermate controls. Similar to the previous experiment, 3 mg/kg of scopolamine significantly increased distance traveled in WT littermate mice compared to vehicle injected WT littermate control mice (1786.0 + 256.6 cm from 90 – 150 min for vehicle, 8952.0 + 789.8 cm for 3 mg/kg scopolamine; One-way ANOVA with Tukey’s Multiple Comparison post-hoc test; F_5,72_=26.7; Figure 1C, E; p < 0.001). In M_1_ knockout animals, 3 mg/kg scopolamine elicited a hyperlocomotor response as compared to vehicle treated M_1_ KO mice (2093.0 + 388.1 cm for vehicle treated M_1_ KO, 7357.0 + 1427.0 cm for M_1_ KO after 3 mg/kg scopolamine; Figure 1C, E p < 0.001). In contrast, the scopolamine-induced increase in locomotor activity observed in WT mice was largely and significantly absent in M_4_ global KO animals (1492.0 + 248.7 cm for vehicle treated M_4_ KO mice, 3574.0 + 407.3 cm for M_4_ KO administered 3 mg/kg scopolamine; Figure 1D, E p > 0.05). When compared to WT animals dosed with 3 mg/kg of scopolamine, M_4_ KO mice administered scopolamine had a significantly reduced distance traveled from 90 – 150 min (Figure 1E, p < 0.001). This suggests that M_4_ plays a dominant role in mediating the scopolamine-induced increase in locomotor activity.

**Figure 1.**
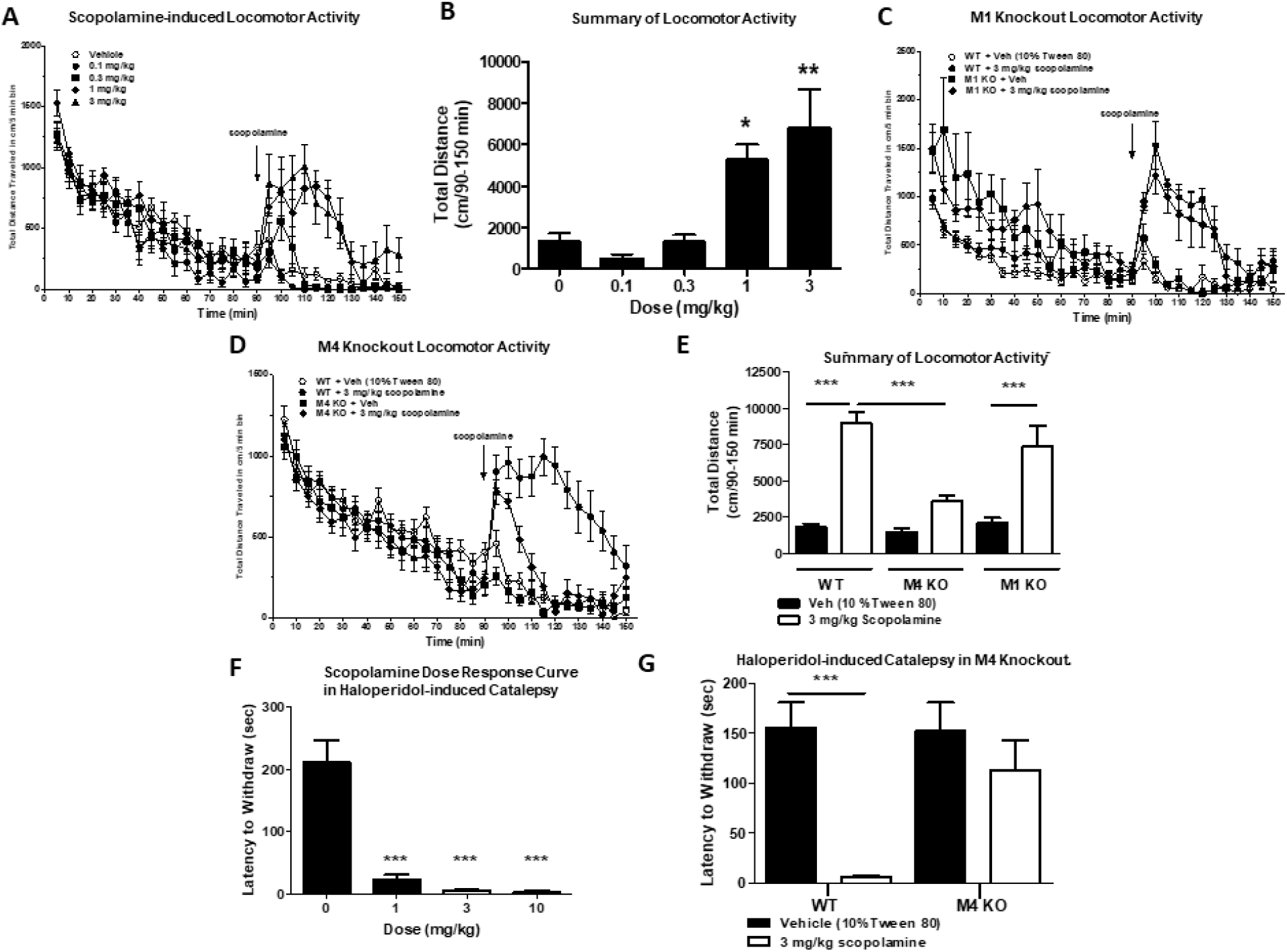
M_4_ underlies the anti-parkinsonian and locomotor-stimulating effects of the muscarinic acetylcholine antagonist scopolamine. Scopolamine induces a dose-dependent increase in locomotor activity (A, B), which persists in M_1_ knockout mice (C, E) but is largely absent in M_4_ knockout mice (D, E). Scopolamine significantly reverses haloperidol-induced catalepsy in wildtype mice, an effect that is absent in M_4_ knockout mice. One-way ANOVA with Dunnett’s or Tukey’s Multiple Comparison post-hoc test; * p < 0.05, ** p < 0.01, *** p < 0.001.

We also examined if M_4_ was responsible for the anti-parkinsonian efficacy of scopolamine in a model of parkinsonism with predictive validity for anti-parkinsonian efficacy, haloperidol-induced catalepsy (HIC)^24^. Similar to the locomotor assay, we first performed a dose-response study to find the dose of scopolamine that was maximally efficacious in reducing catalepsy. Two hours after injection with haloperidol (1 mg/kg, 10 ml/kg, 0.25% lactic acid in water, i.p.) and a dose of scopolamine (15 min before testing, 1 – 10 mg/kg, 10 ml/kg, 10% Tween 80, i.p.), the latency of WT mice to remove their forepaws from an elevated bar was assessed. Scopolamine induced a dose-dependent reversal of HIC (Figure 1F, One-way ANOVA with Dunnett’s post-hoc test; F_3,27_=34.8, p < 0.0001, Figure 1F). In vehicle-treated mice the mean latency to withdraw was 210.9 + 35.9 seconds. The administration of scopolamine at all doses tested significantly reduced the mean latency to withdraw in these mice, and the maximally efficacious dose was again 3 mg/kg (23.4 + 6.7 seconds for 1 mg/kg, 4.4 + 2.3 seconds for 3 mg/kg, 2.9 + 1.8 seconds for 10 mg/kg; Figure 1F, *** p < 0.0001). This dose of scopolamine was then used in the same assay in global M_4_ KO mice and WT littermate controls. In WT littermate controls, 3 mg/kg scopolamine significantly reversed catalepsy (155.9 + 25.3 seconds for vehicle, 6.1 + 1.4 seconds for 3 mg/kg scopolamine; One-way ANOVA with Dunnett’s; F_3,34_=8.7 p=0.0002; Figure 1G ***p<0.001). Vehicle treated M_4_ global KO mice demonstrated similar cataleptic behavior to vehicle treated WT littermates (151.9 + 28.9 seconds for M_4_ KO vs 155.9 + 25.3 seconds for WT vehicle; Figure 1G p>0.05). The administration of 3 mg/kg of scopolamine to M_4_ KO animals did not significantly reduce the mean latency to withdraw compared to WT vehicle treated mice (113 + 29.5 seconds for M_4_ KO; p < 0.05, Figure 1G). Taken together these data indicate that M_4_ is the primary mAChR responsible for the anti-parkinsonian efficacy of the non-selective anti-mAChR antagonist, scopolamine, in HIC.

### Discovery and synthesis of first-in-class M_4_ selective antagonists

Our data implicating M_4_ as the primary mAChR subtype responsible for the locomotor-stimulating and anti-parkinsonian effects of scopolamine gave clear rationale for the development of the first truly selective M_4_ antagonist tool compounds. Utilizing a recently reported partially selective anti-muscarinic compound PCS1055,^25^ a human M_4_ preferring antagonist with potent acetylcholinesterase activity, as a starting point, we devoted significant medicinal chemistry efforts to discover a novel, first-in-class series of selective M_4_ antagonists. Here we reported novel, M_4_ antagonists with balanced activity at both human and rat M_4_ receptors that were devoid of acetylcholinesterase activity and engendered *in vitro* and *in vivo* drug metabolism and pharmacokinetic (DMPK) profiles suitable for *in vivo* proof of concept studies to be performed. Final compounds were prepared as described in Scheme 1, and structures were reported in Figure 2. Briefly, *exo*-amine (**1**) (prepared as previously reported^26^) underwent nucleophilic aromatic substitution (SNAr) with 3,6-dichloropyridazine (**2**) to give chloropyridazine **3**. Boc-deprotection, followed by reductive amination with tetrahydro-2H-pyran-4-carbaldehyde (**4**) gave **5**, which was then substituted via either Suzuki-Miyaura coupling to give **6** (VU6013720) and **7** (VU6021625), or SNAr to give **8** (VU6021302). Analogs can also be synthesized by first substituting chloropyridazine **5** (under identical conditions for the generation of **6-8**), followed by Boc-deprotection and reductive amination. Detailed methods, ^1^H-NMR, and ^13^C-NMR for each final compound (**6-8**) and intermediates were described in the supplemental information.

**Figure 2.**
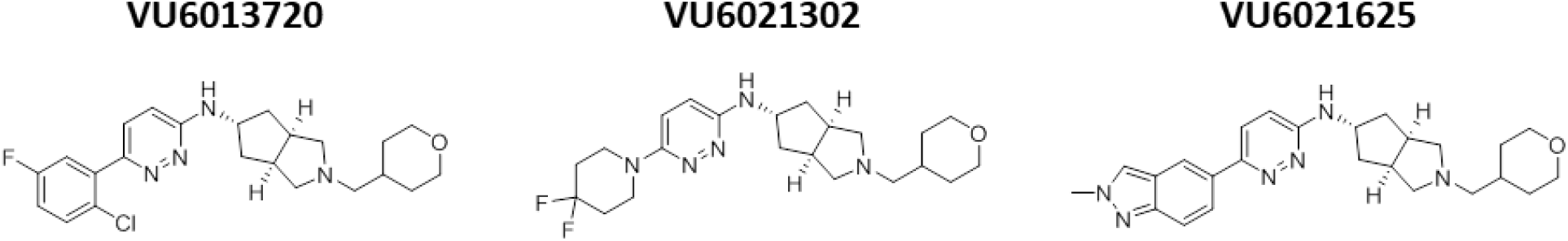
Structure of highly optimized M_4_ antagonists. Chemical structures of novel M_4_ antagonists.

### Pharmacological Characterization of VU6013720, VU6021302, and VU6021625

To understand the potency and selectivity of our novel series of compounds, we tested VU6013720, VU6021302, and VU6021625 in a calcium mobilization assay in cell lines that express individual mAChR receptors^27,28^. First, we examined the potency of each compound in blocking the effects of an EC_80_ concentration of ACh, using calcium mobilization in CHO cell lines that express either rat or human M_4_ as a readout. Each of the novel M_4_ antagonists inhibited the response to ACh with the IC_50_ values listed for VU6013720 (rM_4_ IC_50_ = 20 nM, hM_4_ IC_50_ = 0.59 nM), VU6021302 (rM_4_ IC_50_ = 70 nM, hM_4_ IC_50_ = 1.8 nM), and VU6021625 (rM_4_ IC_50_ = 57 nM, hM_4_ IC_50_ = 0.44 nM) (Figure 3A-C and Table 1). Interestingly, this series of compounds is more potent at human M_4_ than rat M_4_.

**Figure 3.**
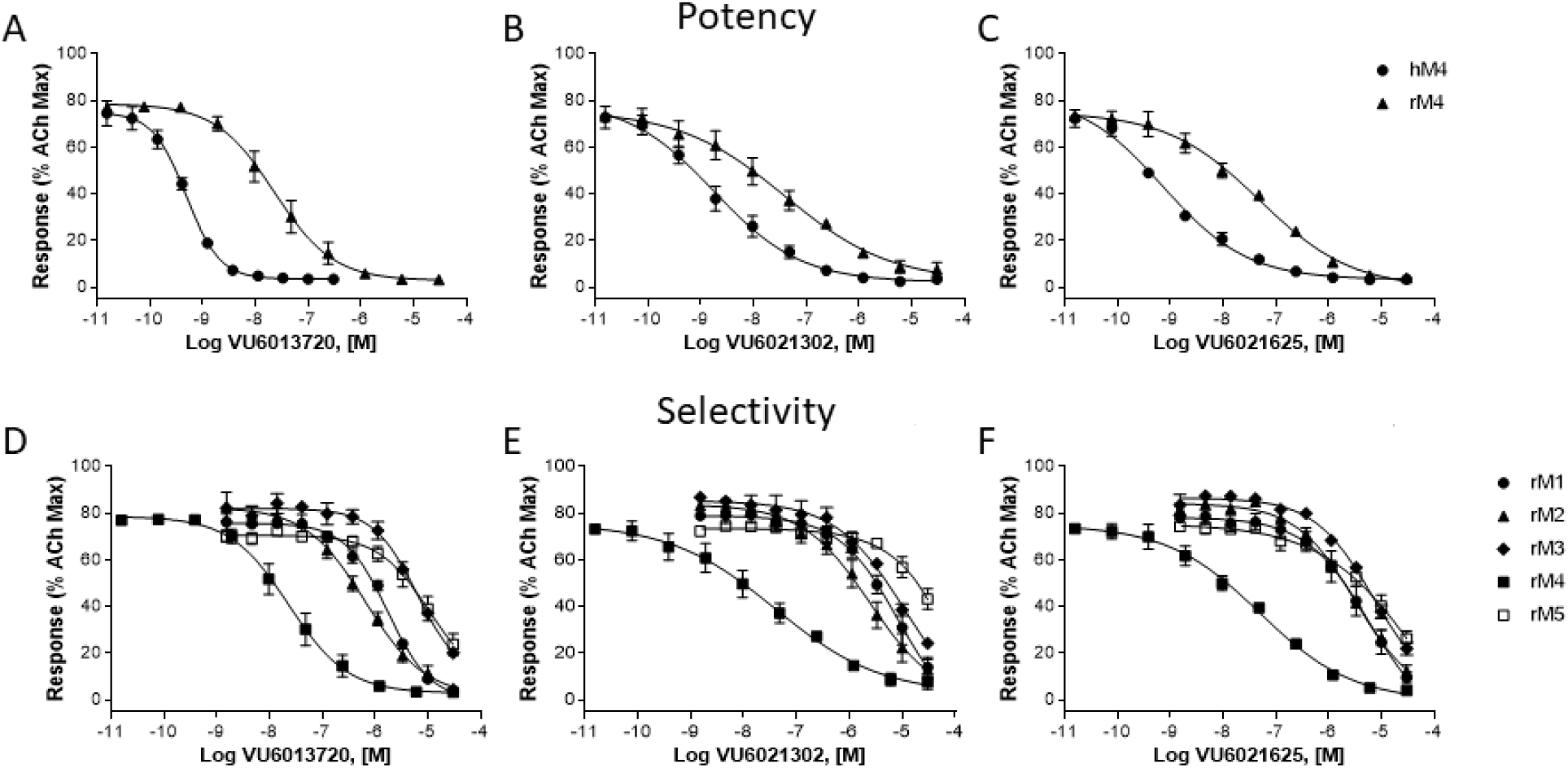
Potency and selectivity of VU6013720, VU6021302, and VU6021625. Potencies were determined by adding a concentration-response curve of M_4_ antagonist followed by an EC_80_ of acetylcholine in human, rat or mouse M_4_-expressing CHO cells. VU6013720, VU6021302, and VU6021625 induced a concentration-dependent inhibition of the release of calcium (A – C). Selectivity of these M_4_ antagonists was evaluated by adding a concentration-response curve of compound followed by an EC_80_ of acetylcholine in M_1_, M_2_, M_3_, M_4_ or M_5_ expressing CHO cells. Data represent the mean ± S.E.M. of 3 independent experiments performed in duplicate.

**Table 1.** Fold Selectivity of VU6013720, VU6021302, and VU6021625. Fold selectivity of M_4_ antagonists over other muscarinic acetylcholine receptors and summary of IC_50_ values.

| Compound |  | human M<sub>4</sub> | rat M<sub>4</sub> | rat M<sub>1</sub> | rat M<sub>2</sub> | rat M<sub>3</sub> | rat M<sub>5</sub> |
| --- | --- | --- | --- | --- | --- | --- | --- |
| VU6013720 | IC<sub>50</sub> (nM) | 0.59 | 20 | 1700 | 670 | > 10,000 | > 10,000 |
|  | [pIC<sub>50</sub> ± SEM] | [9.23 ± 0.04] | [7.71 ± 0.19] | [5.76 ± 0.04] | [6.17 ± 0.04] |  |  |
|  | fold selectivity |  |  | 85 | 34 | > 100 | > 100 |
| VU6021302 | IC<sub>50</sub> (nM) | 1.8 | 70 | >10,000 | 2500 | > 10,000 | > 10,000 |
|  | [pIC<sub>50</sub> ± SEM] | [8.75 ± 0.05] | [7.15 ± 0.10] |  | [5.61 ± 0.10] |  |  |
|  | fold selectivity |  |  | >100 | 36 | > 100 | > 100 |
| VU6021625 | IC<sub>50</sub> (nM) | 0.44 | 57 | 5500 | 3200 | > 10,000 | > 10,000 |
|  | [pIC<sub>50</sub> ± SEM] | [9.36 ± 0.13] | [7.25 ± 0.06] | [5.26 ± 0.16] | [5.50 ± 0.02] |  |  |
|  | fold selectivity |  |  | 96 | 56 | > 100 | > 100 |

We then repeated these calcium mobilization assays in cell lines that express the rat M_1_, M_2_, M_3_ or M_5_ receptors to examine the selectivity of these compounds relative to other mAChR subtypes. VU6013720, VU6021302, and VU6021625 all show functional selectivity at rM_3_ and rM_5_ with IC_50_ values >10,000 nM and selectivity over M_4_ of >100 (Figure 3D-F and Table 1). At rM_1_, VU6013720 has an IC_50_ = 1700 nM with 85 fold selectivity, VU6021302 has an IC_50_ >10,000 nM with >100 fold selectivity, and VU6021625 has an IC_50_ = 5500 nM with 96 fold selectivity (Figure 3D-F and Table 1). The greatest challenge for this series of compounds was selectivity with regard to M_2_. At rat M_2_, VU6013720 has an IC_50_ = 670 nM with 34 fold selectivity, VU6021302 has an IC_50_ = 2500 nM with 36 fold selectivity, and VU6021625 has an IC_50_ = 3200 nM with 56 fold selectivity over r M_4_ (Figure 3D-F and Table 1).

Due to the overall selectivity, especially with regard to M_2_, and potency profile of VU6021625, we chose this as our lead tool compound and submitted this compound for ancillary pharmacology profiling. We examined VU6021625 in the Eurofins radioligand binding panel that utilizes 78 separate G protein-coupled receptors (GPCRs), ion channels, and transporters. With this assay, which screened VU6021625 binding to this panel of targets at 10 μM, we observed little off target binding (see Supplementary Table 1). Outside of mAChRs, only the Histamine H3 receptor showed appreciable binding (88% radioligand displacement), and modest binding to nicotinic α_3_β_4_ and serotonin 5-HT_2B_ receptor was also observed (55% and 53% displacement respectively, Supplementary Table 1). With this overall potency, selectivity, and specificity profile, VU6021625 represents an excellent first-in-class tool compound to examine the effects of M_4_ antagonism.

### VU6021625 *ex vivo* blocks mAChR-induced decreases in D_1_-SPNs activity, DA release, and DA signaling

Previously, using M_4_ positive allosteric modulators and global or conditional M_4_ KO animals, we have shown using *ex* vivo electrophysiology that M_4_ can tonically inhibit the BG direct pathway/D_1_-SPNs as well as cause a sustained inhibition of DA release^15,19^. Using our new tool M_4_ antagonist compound VU6021625, we examined its ability to reverse electrophysiological measures of M_4_ inhibition of direct pathway activity and DA release. First, as previously described, we used whole-cell electrophysiology to patch into GABAergic cells of the SNr and recorded miniature inhibitory post-synaptic currents (mIPSCs) to examine direct pathway neurotransmitter release probability^15^. After recording a baseline period of 5 minutes, we bath applied 10 μM VU6021625 and recorded mIPSC frequency. Inhibition of M_4_ activity in the direct pathway increased mIPSC frequency by ~40%, indicating that M_4_ tonically inhibits the BG direct pathway, and that removal of this inhibition increases direct pathway output, as would be predicted by our previous work (Figure 4A-B, paired t-test, *p < 0.05). To further evaluate effects of VU6021625 on mAChR mediated inhibition of transmission from D_1_-SPN terminals, we again recorded mIPSCs but bath applied 10 μM of the non-selective mAChR agonist oxotremorine-M (Oxo-M), which decreased D_1_-SPN mIPSC frequency by ~20% (Figure 4C-D). We have preciously shown that the effects of Oxo-M at this synapse are mediated exclusively by M_4_^15^. Bath application of VU6021625 completely blocked this effect, indicating that M_4_ antagonism can block tonic activation of M_4_ by endogenous ACh as well as M_4_-mediated effects on inhibitory transmission in the direct pathway that are induced by exogenously added agonists (Figure 4C-D, paired t-test, **p<0.01).

**Figure 4.**
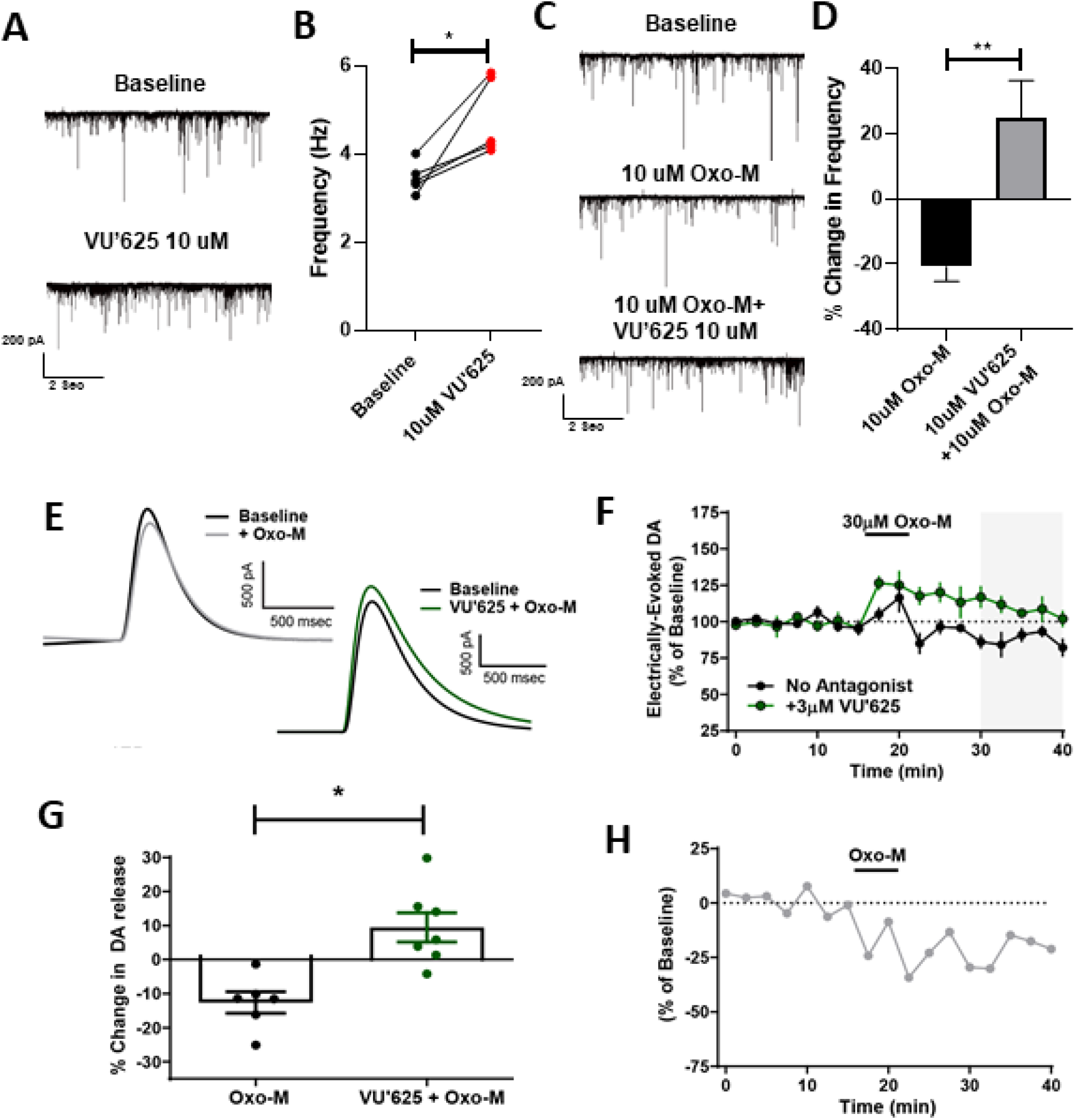
VU6021625 reversed muscarinic induced deficits in dopamine release and signaling. Sample traces of miniature inhibitory post-synaptic currents (mIPSCs) during baseline (top) and following bath application of 10 μM VU6021625 (bottom) (A). Inhibition of M_4_ activity by VU6021625 significantly increased mIPSC frequency (paired t-test, *p<0.05) (B). Sample traces of mIPSCs during baseline (top), bath application of the non-selective mAChR agonist oxotremorine-M (Oxo-M) (middle), or bath application of 10 μM Oxo-M + VU6021625 (bottom) (C). Bath application of VU6021625, completely blocked the change in frequency induced by Oxo-M, paired t-test, **p<0.01 (C-D). Sample traces (E) and time-courses (F) of Oxo-M-induced inhibition of DA release in the absence or presence of the M_4_-selective antagonist VU6021625. Bar graph summary depicting the % inhibition of DA release observed under different conditions from 30-40 min (n=6-7, *: p < 0.005, two tailed Mann Whitney test) (G). Time-course of M_4_-mediated effects obtained from subtracting the mean values for the time-course in absence of antagonist from the mean values of the time-course in the presence of VU6021625 (H).

### Application of the M_4_-selective antagonist VU6021625 unmasks a robust Oxo-M-dependent increase in DA release

Beyond the effects of M_4_ inhibition of direct pathway activity, we have also previously shown that M_4_ potentiation causes a sustained inhibition of DA release in the dorsal striatum^19^. To determine the ability of the M_4_ receptor to modulate striatal DA release we monitored Oxo-M mediated changes in electrically-evoked DA via fast-scan cyclic voltammetry in the dorsolateral striatum in the absence or presence of VU6021625 (Figure 4E-G). All experiments reported here were performed in the presence of nicotinic acetylcholine receptor (nAChR) antagonist (DhβE; 1μM) to remove nAChR-mediated DA release. In the absence of M_4_ antagonism, we saw a transient increase in DA when Oxo-M was applied followed by a sustained inhibition after drug washout (sustained inhibition of −12.64 ± 3.18 % of baseline DA release). Inclusion of VU6021625 (3μM) caused an increase in the amount of DA release (Figure 4F) and significantly reversed the direction of Oxo-M effects on DA release observed after Oxo-M washout resulting in a net increase in DA release (sustained increase of 9.45 ± 4.28 % of baseline DA release; Figure 4G). To determine the specific contribution of M_4_ receptor activation on DA release we subtracted the mean values of the Oxo-M time-course obtained in the absence of any antagonist from the mean values obtained in the presence of VU6021625 (Figure 4H), which revealed a rapid onset of M_4_-mediated inhibition of DA release that was sustained at time-points well after Oxo-M has washed out of the bath.

The results obtained with VU6021625 in fast-scan cyclic voltammetry support our previous finding that M_4_ receptor activation can induce a sustained inhibition of DA release in the dorsolateral striatum in the presence of nAChR antagonists^17^. Unlike our previous report where Oxo-M induced a suppression at all time-points examined, we found that 30 μM Oxo-M alone caused a transient increase in DA release followed by a sustained inhibition. However, in the presence of VU6021625, an Oxo-M induced increase in DA release was observed at all time-points (Figure 4). While this is consistent with M_4_ receptor activation leading to a sustained inhibition of DA release, the identity of the mAChR mediating this increase in DA release is still not know. Studies in the nucleus accumbens have shown that M_5_ receptors can mediate an enhancement of DA release in this brain area raising the possibility that M_5_ receptors may also be able to increase DA release in the dorsolateral striatum. These results suggest that there could be a competition between different mAChR-subtypes in the striatum with activation of some subtypes leading to inhibition of DA release while activation of other mAChR subtypes could lead to a potentiation. Future studies will be needed to gain critical insights into what mAChR subtypes mediate these increases in DA release in the dorsolateral striatum and elucidate if there are physiological conditions that favor the activation of DA inhibiting mAChRs and DA potentiating mAChRs.

### VU6021625 possesses DMPK properties suitable for *in vivo* use

Before use in animal models of parkinsonism and dystonia, we assessed the DMPK properties of VU6021625 to determine if VU6021625 was suitable for *in vivo* use in rodents and to inform the dosing paradigms for behavioral studies. Mouse plasma and brain pharmacokinetics (PK) were obtained over a 7 hr time course following a single 1 mg/kg i.p. dose (vehicle: 20% β-cyclodextrin 80% water [w/v]; 10 mL/kg body weight) to male, C57Bl/6 mice (*n* = 3 per time point) (Figure 5A). The maximum total concentration of the compound in plasma (C_max,total_) was 170 ng/mL (393 nM) with a time-to-reach C_max_ (T_max_) of 0.25 hr. In brain, a C_max,total_ of 31.6 ng/mL (73.0 nM) was observed with a T_max_ of 1 hr. Distribution to brain from plasma on an area-under-the-curve (AUC)-based total concentration basis (K_p_) was moderate (0.25, Figure 5A).

**Figure 5.**
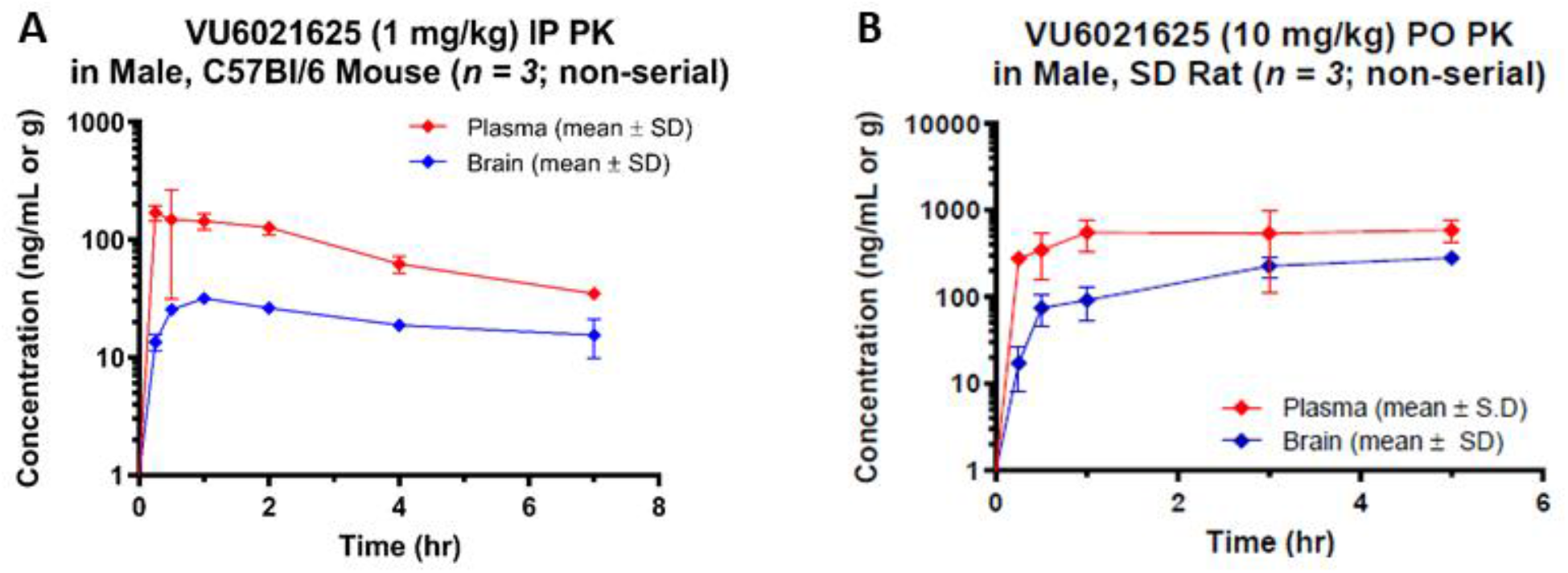
*In Vivo* Pharmacokinetics of VU6021625. Plasma and brain concentrations of VU6021625 were measured following systemic administration (1 mg/kg, 10 ml/kg, i.p.) in mice (A) and (10 mg/kg, 10 ml/kg, p.o.) in rats (B). SD = Sprague Dawley

Rat plasma and brain PK were obtained over a 5 hr time course following a single 10 mg/kg per oral (p.o.) dose (vehicle: 0.5% methylcellulose 95.5% water [w/v]; 10 mL/kg body weight) to male, Sprague Dawley rats (*n* = 3 per time point) (Figure 5B). The total C_max_ in plasma was 586 ng/mL (1350 nM) with a T_max_ of 5 hr. The total C_max_ in brain was 282 ng/mL (652 nM) with a T_max_ of 5 hr. AUC-based total brain distribution was moderate (K_p_ = 0.35) while unbound distribution was low (K_p,uu_ = 0.13) based on *in vitro* rat plasma and brain homogenate binding data (fu_plasma_ = 0.563, fu_brain_ = 0.206) from equilibrium dialysis assays.

In light of the low brain:plasma K_p,uu_ observed in the rat study, potential P-glycoprotein (P-gp) substrate activity of VU6021625 was evaluated using an *in vitro* bidirectional efflux assay with MDCK-MDR1 cells (performed via contract by Absorption Systems LLC [Exton, PA]). This experiment revealed a high efflux ratio (ER) of 71 indicating that the compound is subject to P-gp-mediated active efflux (Table 2). However, an absolute unbound brain concentration (134 nM) > 2-fold higher than the compound’s *in vitro* rat M_4_ potency was achieved in the rat PK study thereby demonstrating its *in vivo* utility despite P-gp efflux.

**Table 2.** Pharamacokinetic Properties of VU6021625. Summary of *in vivo* and *in vitro* characteristics of VU6021625

| Property | Value |
| --- | --- |
| MDCK-MDR1 (Efflux Ratio) | 71 |
| $f_u$ brain (r) | 0.206 |
| $f_u$ plasma (r) | 0.563 |
| Kinetic Solubility (μM) | 95.1 ± 10.3 (pH 2.2)<br>90.8 ± 9.62 (pH 6.8) |

### VU6021625 has anti-parkinsonian efficacy

To test if VU6021625 has anti-parkinsonian efficacy as expected from our data presented in Figure 1, we utilized the HIC animal model of parkinsonian motor deficits. Vehicle treated mice displayed a mean latency to withdraw their forepaws of 202.4 + 16.6 seconds. Administration of 0.3 mg/kg of VU6021625 did not significantly reduce catalepsy in these mice (159.0 + 27.3 seconds, Figure 6A, B, 10 ml/kg, 20% (2-Hydroxypropyl)-Beta-Cyclodextrin (HPBCD), i.p; One-way ANOVA with Dunnett’s post-hoc test F_3,44_=8.6, p>0.05). However, administration of 1 or 3 mg/kg significantly reversed cataleptic behavior when compared to vehicle treated mice (79.6 + 20.2 seconds for 1 mg/kg, 74.3 + 19.5 for 3 mg/kg Figure 6A, B,; 10 ml/kg, 20% HPBCD, i.p; One-way ANOVA with Dunnett’s post-hoc test F_3,44_=8.6 p < 0.0001).

**Figure 6.**
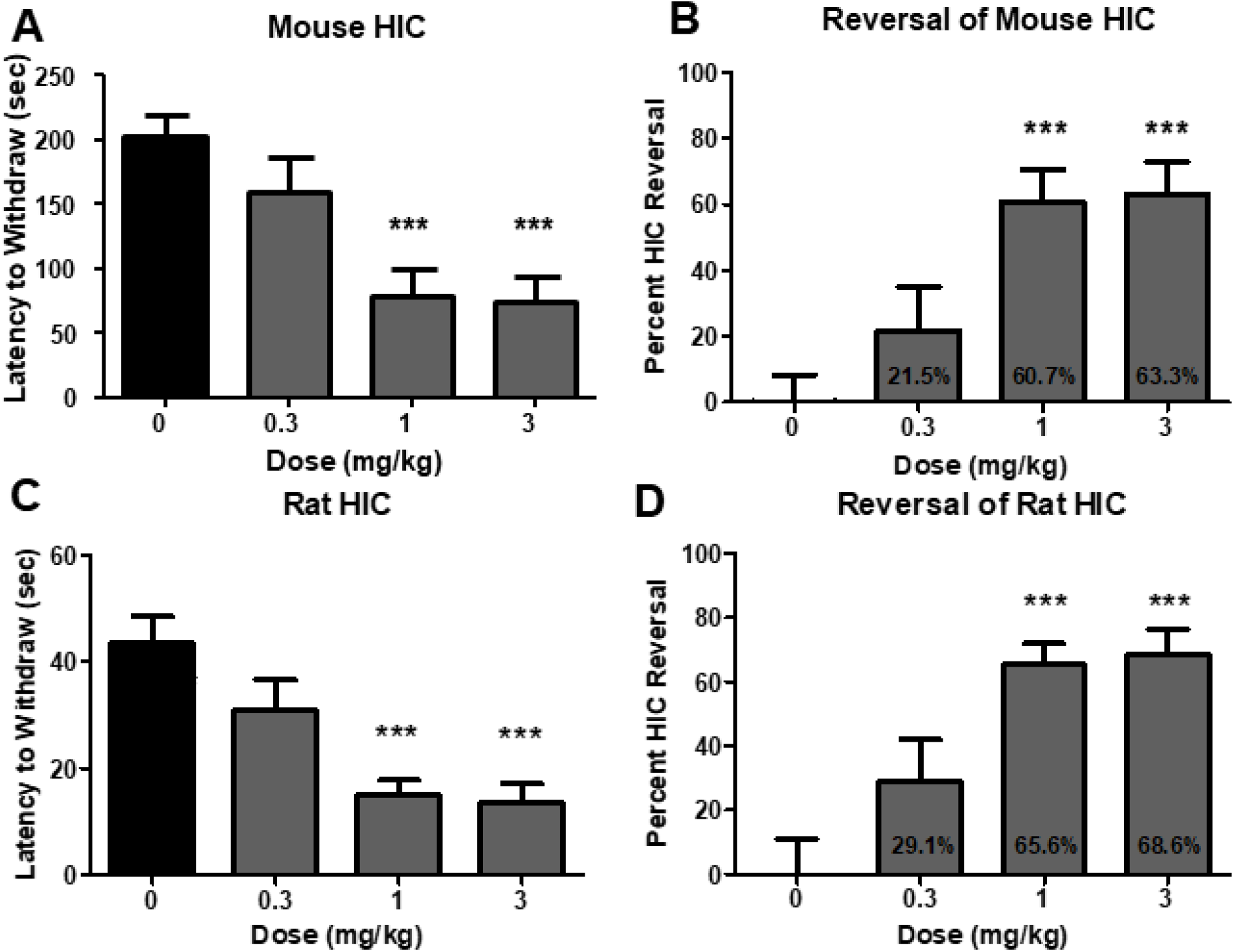
VU6021625 displays anti-parkinsonian efficacy. Systemic administration of VU6021625 demonstrates dose-dependent efficacy in the haloperidol-induced catalepsy animal model of parkinsonian motor deficits in both mice (A, B) and rats (C, D) with a minimal effective dose of 1 mg/kg in both species. One-way ANOVA with Dunnett’s post-hoc test; *** p < 0.001.

In order to demonstrate that this was not a species-specific effect, VU6021625 was also tested in a rat HIC model. In rats, VU6021625 reduced the cataleptic phenotype in this model at the same doses needed for efficacy in mice. Vehicle treated rats exhibited a mean latency to withdraw of 43.6 + 4.8 seconds. The administration of 0.3 mg/kg of VU6021625 did not significantly reduce mean latency to withdraw in these rats (30.9 + 5.7 seconds, Figure 6C, D, 1 ml/kg, 10% Tween 80/0.5% methylcellulose, i.p.; One-way ANOVA with Dunnett’s post-hoc test, F_3,36_= 10.9, p < 0.05). The administration of 1 or 3 mg/kg significantly reversed cataleptic behavior when compared to vehicle treated rats (15.0 + 2.7 seconds for 1 mg/kg, 13.7 + 3.3 for 3 mg/kg; Figure 6C, D 1 ml/kg, 10% Tween 80/0.5% methylcellulose, i.p.; One-way ANOVA with Dunnett’s post-hoc test, F_3,36_=10.9, ***p < 0.001). This equated to a significant reversal of catalepsy with 1 mg/kg or 3 mg/kg of VU3021625 (29.1 + 13.0% reversal for 0.3 mg/kg, 65.6 + 6.2% reversal for 1 mg/kg, 68.8 + 7.7% reversal for 3 mg/kg; One-way ANOVA with Dunnett’s post-hoc test, Figure 6D ***p < 0.001). The minimum effective dose for VU6021625 in this assay was also 1 mg/kg. Additionally, we collected plasma and brain samples at the end of the rat behavioral study to determine the concentration of VU6021625 by LC-MS/MS analysis. At the 1 and 3 mg/kg dose levels, the compound reached unbound brain concentrations of 12.3 nM and 34.2 nM, respectively (Table 3). These concentrations align well with the potency of this compound at rat M_4_ and are well below the potency of the other four mAChR subtypes (Table 1), and indicate that VU6021625 has anti-parkinsonian efficacy in mouse and rat models at exposure levels specific for M_4_.

**Table 3.** Pharamacokinetic-Pharmacodynamic relationship of VU6021625. Unbound brain drug concentrations of VU6021625 after catalepsy testing. At the minimally efficacious dose (1mg/kg) drug exposure reaches levels that are consistent with selectivity *in vitro*.

| **VU6021625 HIC Dose**<br>**Response** |  |
| --- | --- |
| **Dose**<br>**(mg/kg)** | Brain<br>Unbound (nM) |
| **0.3** | 4.8 |
| **1** | 12.3 |
| **3** | 34.2 |

### VU6021625 has anti-dystonic efficacy

Trihexyphenidyl, a non-selective mAChR antagonist, is the only oral medication for dystonia that has been proven effective in a double-blind, placebo controlled clinical trial^11,12^. To determine if VU6021625 is also effective, we tested a genetic model of DOPA-responsive dystonia (DRD) that carries a point mutation in the mouse tyrosine hydroxylase gene that diminishes DA synthesis and release, and is synonymous with human forms of the disease^29^. The abnormal dystonic movements in DRD mice are ameliorated by trihexyphenidyl, similar to human patients^29^. Administration of VU6021625 (1-3 mg/kg, 10 ml/kg, 1% Tween80, i.p.) significantly reduced the dystonic movements observed in DRD mice at the 3 mg/kg dose (Figure 7, repeated measures ANOVA, F3,21=4.85, Dunnett’s multiple comparisons post hoc test, *p < 0.05), providing similar efficacy to trihexyphenidyl (paired t-test, ***p < 0.001). These data demonstrate that VU6021625 has anti-dystonic efficacy in this mouse model, as well as anti-parkinsonian efficacy.

**Figure 7.**
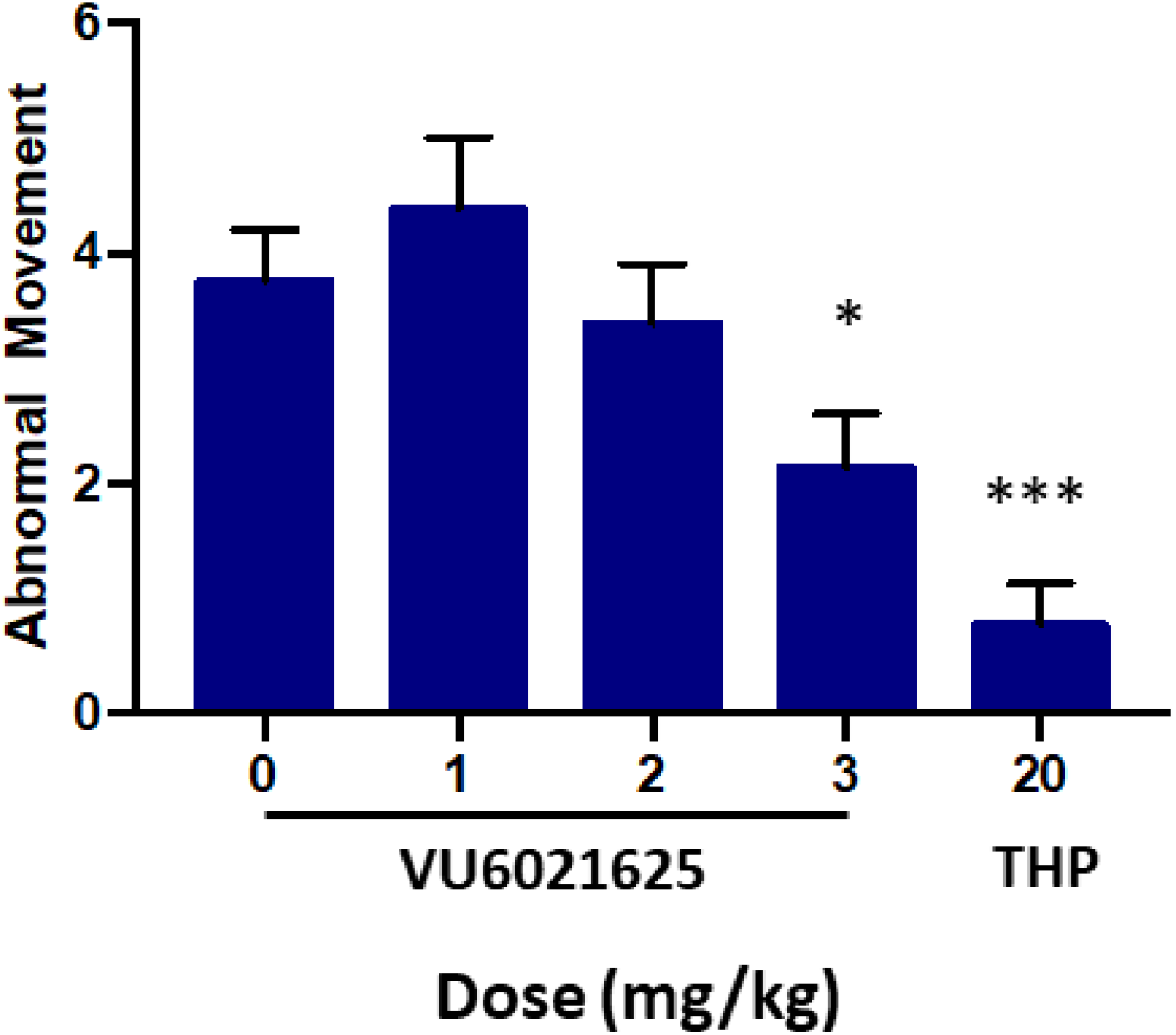
VU6021625 displayed anti-dystonic efficacy. Using a genetic model of DOPA-responsive dystonia (DRD) we demonstrated that VU6021625 significantly reduced abnormal movements observed in these mice with a minimal effective dose of 3 mg/kg (1-3 mg/kg, 10 ml/kg, 1% Tween 80, i.p.). Repeated measures ANOVA with Dunnett’s multiple comparisons post hoc test; * p < 0.05), providing similar efficacy to trihexyphenidyl (THP, paired t-test compared to vehicle, *** p < 0.001).

## Discussion

We report the discovery and characterization of a novel class of M_4_ antagonists that mark a major advance in the selectivity and specificity over previously reported compounds. The most potent and selective of these compounds, VU6021625, exhibits the ability to reverse effects of mAChR agonists on BG activity and DA release in mouse brain slices, and exert anti-parkinsonian and anti-dystonic efficacy in pharmacological and genetic animal models of movement disorders. Importantly, in these models we have established a PK-PD relationship that demonstrates that at doses needed for efficacy, unbound brain concentrations of VU6021625 are well within the selectivity range that VU6021625 possesses over other mAChRs. These findings, as well as our data utilizing genetic KO mice of M_4_ with non-selective compounds, implicate M_4_ as the primary mAChR subtype that mediates the locomotor-stimulating, anti-dystonic, and anti-parkinsonian effects of non-selective anti-mAChR antagonists, although it is possible that other mAChRs, namely M_1_, may also play a modulatory role to muscarinic anti-parkinsonian efficacy as well^30^.

The ability of VU6021625 to reverse the cataleptic phenotype in HIC studies suggests that M_4_ selective antagonists may be beneficial in removing hypokinetic aspects of parkinsonian motor deficits, such as bradykinesia and rigidity. Testing the role of M_4_ selective antagonists in other rodent models as well as other aspects of parkinsonian motor impairments including tremor and gait disturbances will be key to understanding the extent of anti-parkinsonian effects of M_4_ antagonists. Non-selective mAChR antagonists have also shown efficacy in reducing the prevalence of some of these other parkinsonian motor symptoms. Mechanistic determination of the specific role of M_4_ antagonism in the efficacy of these non-selective compounds in treating other parkinsonian symptoms will be critical to understanding the potential utility of M_4_ antagonists as a monotherapy in PD. For example, M_4_ potentiation using VU0467154 has been previously shown to relieve dyskinesia induced by chronic L-DOPA administration^31^, suggesting that in treatment-induced abnormal movements in PD, M_4_ activity may be low in the dyskinetic state^9^. Whether M_4_ antagonists would further exacerbate dyskinesia, or if repeated administration of M_4_ antagonists could cause dyskinesia remains to be tested. A greater understanding of the ability of M_4_ to cause or alter abnormal movements will be important to understanding the full utility of M_4_ antagonists in the treatment of movement disorders.

While the etiology of dystonia is complex^5^, a number of genes have been shown to be causative for dystonia^32^. Some of these genes, such as the genes which are causative for DRD, have clear links to the synthesis, degradation, or signaling of DA^6,29,33^. However, several genetic forms of dystonia, such as *Dyt1*, have no clear link to DA, although some mouse models of these genetic forms of dystonia do show DA deficits^34,35^. Similar to trihexyphenidyl, which is used clinically to treat movement disorders, VU6021625 reduces abnormal movements in one of these DA-focused models of dystonia. Whether VU6021625 can relieve behavioral and pathological correlates of dystonia in other forms of genetic dystonia not linked to DA, as well as in other DA linked models, has yet to be determined but will be critical to our understanding of the range of utility of M_4_ antagonism in dystonia as well as mechanistically understanding the etiology of different forms of dystonia.

VU6021625 represents a major advance in the discovery of selective M_4_ antagonists, and provides, for the first time, an M_4_ antagonist tool compound that can be used to explore the role M_4_ plays in movement disorders *in vivo*. Our initial data support the hypothesis that M_4_ blockade may underlie a majority of the efficacy observed with non-selective anti-mAChR therapeutics in movement disorders. The development of this *in vivo* tool compound enables a number of studies that will help understand the role of M_4_ in the treatment and etiology of several movement disorders. These data will provide critical pre-clinical rationale for the further development and optimization of M_4_ antagonists and selective blockade of M_4_ represents a potential novel treatment mechanism to meet the huge unmet clinical need across several movement disorders.

## Methods

### General Chemistry Methods

All reactions were carried out employing standard chemical techniques under inert atmosphere. Solvents used for extraction, washing, and chromatography were HPLC grade. All reagents were purchased from commercial sources and were used without further purification. Low resolution mass spectra were obtained on an Agilent 6120 or 6150 with UV detection at 215 nm and 254 nm along with ELSD detection and electrospray ionization, with all final compounds showing >95% purity and a parent mass ion consistent with the desired structure. All NMR spectra were recorded on a 400 MHz Brüker AV-400 instrument. ^1^H chemical shifts are reported as δ values in ppm relative to the residual solvent peak (MeOD = 3.31). Data are reported as follows: chemical shift, multiplicity (br = broad, s = singlet, d = doublet, t = triplet, q = quartet, p = pentet, dd = doublet of doublets, ddd = doublet of doublet of doublets, td = triplet of doublets, dt = doublet of triplets, m = multiplet), coupling constant, and integration. ^13^C chemical shifts are reported as δ values in ppm relative to the residual solvent peak (MeOD = 49.0). Automated flash column chromatography was performed on a Biotage Isolera 1 and a Teledyne ISCO Combi-Flash system. Microwave synthesis was performed in a Biotage Initiator microwave synthesis reactor. RP-HPLC purification of final compounds was performed on a Gilson preparative LC system. For detailed instructions on synthesis of each compound, see the supplemental information

### Calcium Mobilization Assays

Compound-evoked decreases to an EC_80_ concentration of ACh in intracellular calcium were measured using Chinese hamster ovary (CHO) cells stably expressing mouse, rat, or human muscarinic receptors (M_1_–M_5_; M_2_ and M_4_ cells were co-transfected with G_qi5_). Cell culture reagents were purchased from Gibco-ThermoFisher Scientific (Waltham, MA) unless otherwise noted. Cells (15,000 cells/20 μL/well) were plated in Greiner black wall / clear bottom 384 well plates in F_12_ medium containing 10% FBS, 20 mM 2-[4-(2-hydroxyethyl)piperazin-1-yl]ethanesulfonic acid (HEPES), and 1X Antibiotic/Antimycotic. The cells were grown overnight at 37 °C in the presence of 5% CO_2_. The next day, the medium was removed and replaced with 20 μL of 1 μM Fluo-4, AM (Invitrogen, Carlsbad, CA) prepared as a 2.3 mM stock in dimethyl sulfoxide (DMSO) and mixed in a 1:1 ratio with 10% (w/v) pluronic acid F-127 (Invitrogen, Carlsbad, CA) and diluted in Assay Buffer (Hank’s Balanced Salt Solution (HBSS), 20 mM HEPES, 4.16 mM sodium bicarbonate (Sigma-Aldrich, St. Louis, MO) and 2.5 mM Probenecid (Sigma-Aldrich, St. Louis, MO)) for 50 minutes at room temperature. Dye was removed and replaced with 20 μL of Assay Buffer. Compounds were serially diluted 1:3 into 10 point concentration response curves in DMSO using the AGILENT Bravo Liquid Handler (Atlantic Lab Equipment, Santa Clara, CA), transferred to daughter plates using an Echo acoustic plate reformatter (Labcyte, Sunnyvale, CA) and diluted in Assay Buffer to a 2X final concentration. Ca^2+^ flux was measured using a Functional Drug Screening System 6000 or 7000 (FDSS6000/7000, Hamamatsu, Japan). After establishment of a fluorescence baseline for 2-3 seconds (2-3 images at 1 Hz; excitation, 480 ± 20 nm; emission, 540 ± 30 nm), 20 μL of test compound was added to the cells, and the response was measured. 140 seconds later, 10 μL (5X) of an EC_20_ concentration of ACh (Sigma-Aldrich, St. Louis, MO) was added to the cells, and the response of the cells was measured. Approximately 125 seconds later, an EC_80_ concentration of ACh was added. Calcium fluorescence was recorded as fold over basal fluorescence and raw data were normalized to the maximal response to agonist. Potency (IC_50_) and maximum response (% ACh Max) for compounds was determined using a four-parameter logistical equation using GraphPad Prism (La Jolla, CA) or the Dotmatics software platform:

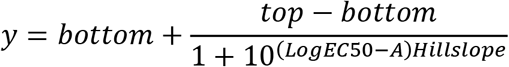

where *A* is the molar concentration of the compound; *bottom* and *top* denote the lower and upper plateaus of the concentration-response curve; HillSlope is the Hill coefficient that describes the steepness of the curve; and EC_50_ is the molar concentration of compound required to generate a response halfway between the *top* and *bottom*.

### Ancillary Pharmacology Screening

VU6021625 was tested at a concentration of 10 μM in the Eurofins Panlabs Lead Profiling Screen, a radioligand binding assay panel consisting of 78 GPCRs, ion channels, and transporters. Displacement of ≥50% radioligand binding at a panel target was considered significant.

### Brain slice preparation

Mice were anesthetized by continuous isoflurane (5%) and then transcardially perfused with a cold solution of 92 mM *N*-methyl-D-glucamine (NMDG), 2.5 mM KCl, 1.2 mM NaH_2_PO_4_, 30 mM NaHCO_3_, 20 mM Hepes, 25 mM D-glucose, 5 mM sodium ascorbate, 2 mM thiourea, 3 mM sodium pyruvate, 10 mM MgSO_4_, 0.5 mM CaCl_2_ (pH 7.3) at 295-300 mOsm. Mice were then decapitated and the brains were removed. 300 μm coronal sections of the SNr or striatum were made on a Leica Vibratome 1200S. After sectioning, coronal slices were submerged for 10–15 min at 32 °C in the same NMDG solution as above. Following this recovery period, slices were transferred to ACSF containing 126 mM NaCl, 2.5 mM KCl, 26.2 mM NaHCO_3_, 1.25 mM NaH_2_PO_4_, 2 mM CaCl_2_, 1.5 mM MgSO_4_, 10 mM D-glucose and 5 mM sodium ascorbate.

### Whole Cell Patch Clamp Electrophysiology

Whole cell voltage-clamp signal was amplified using Axon Multiclamp 700B amplifiers with appropriate electrode-capacitance compensation. Patch pipets were prepared from borosilicate glass using a Sutter instruments P-100 Flaming/Brown micropipet puller, and had resistance of 3–6 MΩ when filled with the following intracellular solution (mM): 130 CsCl, 10 NaCl, 0.25 CaCl_2_, 2 MgCl_2_, 5 EGTA, 10 HEPES, 10 glucose, 2 Mg-ATP. The pH of the pipet solution was adjusted to 7.3 with 1 M CsOH, and osmolarity was adjusted to 285–290. Whole cell recordings were made from visually identified cells in the SNr under an Olympus BX50WI upright microscope (Olympus, Lake Success, NY). A low-power objective (4×) was used to identify the SNr, and a 40× water immersion objective coupled with Hoffman optics was used to visualize the individual neurons. GABAergic cells of the SNr were identified by previously determined membrane characteristics and firing rates^15,36^.

To isolate mIPSCs, slices were perfused continuously at a rate of ~2.0 ml/min with an oxygenated solution containing (in mM): 126 mM NaCl, 2.5 mM KCl, 26.2 mM NaHCO_3_, 1.25 mM NaH_2_PO_4_, 2 mM CaCl_2_, 1.5 mM MgSO_4_, 10 mM D-glucose, pH 7.35 with 0.5 μM tetrodotoxin (TTX), 5 μM AMPA receptor 6-cyano-7-nitroquinoxaline-2,3-dione (CNQX) and 1 μM NMDA receptor antagonist DL-2-amino-4-methyl-5-phosphono-3-pentenoic acid (AP-5). Slices were perfused with this solution at 25 °C for at least 15 minutes following establishment of electrical access. Access resistances were <15 MΩ. mIPSCs were recorded from GABAergic cells of the SNr held at −70 mV in GAP free mode. All drugs were bath-applied with the complete exchange of the external solution not exceeding 30 sec. Data were acquired using Digidata 1440A and pClamp 9.2 and analyzed with Mini Analysis software (Synaptosoft)

### Fast Scan Cyclic Voltammetry

Electrically-evoked DA overflow was monitored with carbon fiber electrodes with a 5 μm diameter as previously described^19^. A triangular voltage wave (−400 to +1000 mV at 300V/sec) was applied to fresh cut carbon fiber electrodes every 100 msec. When monitoring electrically-evoked DA transients, stimulating electrodes were placed 75 μm deep into the dorsolateral striatum and slices were electrically stimulated (30-600 μA) every 2.5 min via a bipolar stimulating electrode placed ~100 μm from the carbon fiber. For typical experiments the stimulation intensity used was 200-400 μA so as to induce both nAChR-dependent and – independent DA release. Current was acquired using a Clampex9.2/Digidata1440A system with a low pass Bessell Filter at 10kHz and digitized at 100kHz. Background-subtracted cyclic voltammograms served both to calibrate the electrodes and to identify dopamine as the substance that was released following electrical stimulation. The best-fit simulation of electrically-evoked dopamine overflow was found by nonlinear regression. All time course data is presented as the mean ± SEM for individual time points. Sustained inhibition was defined as the inhibition that was observed 10-20 minutes after Oxo-M had washed out. These data were analyzed using a two tailed Mann Whitney test and statistical significance was determined as p < 0.05.

### Pharmacokinetics (PK)

The mouse PK study was performed in adult male C57Bl/6 mice (*n* = 3 per time point). In this study, VU6021625 was formulated (0.1 mg/mL) in 20% β-cyclodextrin 80% water (w/v) and administered at 10 mL/kg body weight. Non-serial sampling of plasma and brain was performed at multiple time points (0.25, 0.5, 1, 2, 4, and 7 hr) post-administration. The rat PK study was performed in adult male Sprague-Dawley rats (*n* = 3 per time point). In this study, VU6021625 was formulated (1 mg/mL) in 0.5% methylcellulose 95.5% water (w/v) and administered at 10 mL/kg body weight. Non-serial sampling of plasma and brain was performed at multiple time points (0.25, 0.5, 1, 3, and 5 hr) post-administration. In both studies, brain:plasma K_p_ was determined by division of the brain AUC_0.25-last_ by the plasma AU_C0.25-last_. In the rat study, unbound brain:unbound plasma K_p,uu_ was determined by division of the K_p_ by the [fu_plasma_/fu_brain_] ratio, which was obtained via equilibrium dialysis binding assays with rat plasma (*n* = 1 performed in triplicate) and rat brain homogenate (*n* = 1 performed in triplicate). Plasma and brain homogenate binding assays, bioanalytical sample preparation, and quantitation by LC-MS/MS was performed essentially as described previously^37,39–40^. Data are presented (Figure 5A/B) as means ± SD. Evaluation of VU6021625 for P-gp efflux potential was performed via contract with Absorption Systems LLC (Exton, PA) using a bidirectional transwell assay (*n* = 1 performed in duplicate) with a 5 μM substrate concentration and their standard in-house assay and quantitation procedures.

### Animal Housing

Animals were group-housed under a 12/12 h light-dark cycle with food and water available *ad libitum*. All animal experiments were approved by the Vanderbilt University or Emory University Animal Care and Use Committee, and experimental procedures conformed to guidelines established by the National Research Council *Guide for the Care and Use of Laboratory Animals*.

### Open Field Locomotor Assay

Locomotor activity was tested in wild-type and M_4_ or M_1_ KO mice, 8–12 weeks old, using an open field system (OFA-510, MedAssociates, St. Albans, VT) with three 16 × 16 arrays of infrared photobeams. Scopolamine induced locomotor activity was assessed with the following paradigm: animals were habituated for 90 min in the open field before being injected with vehicle (10% Tween 80, 10 ml/kg, i.p.) or scopolamine (0.1, 0.3, 1, or 3 mg/kg, 10 ml/kg, 10% Tween 80 i.p.), and locomotor activity was recorded for an additional 60 min (150 minute total session length). Data were analyzed using the activity software package (MedAssociates, St. Albans VT) and expressed as total distance traveled in cm per 5 min bin.

### Haloperidol-Induced Catalepsy

Wildtype C57Bl6/J, littermate controls or M_4_ global knockout animals were administered haloperidol (1 mg/kg, 0.25% lactic acid in water, i.p.). For the scopolamine experiments, the mice were administered vehicle (10% Tween80) or scopolamine (i.p.) 105 minutes after haloperidol. For the VU6021625 experiment the animals were administered vehicle (20% HPBCD) or VU6021625 (0.3-3 mg/kg, i.p.) 105 minutes after haloperidol. They were tested 15 minutes later by placing their forelimbs on a raised bar and recording the latency for the animals to remove their forelimbs from the bar with a cutoff of 300 seconds. Adult male Sprague-Dawley rats were injected with 1.5 mg/kg of haloperidol i.p. One hour later the animals were administered 0.3-3 mg/kg of VU6021625, or vehicle. Cataleptic behavior was determined 30 minutes later as previously described^34^ by placing the forelimbs on a bar raised 6 cm above the table and recording the amount of time it takes for the rat to withdraw the forelimbs with a cutoff of 60 seconds. Data are expressed as mean latency to withdraw + SEM or percent inhibition of catalepsy + SEM. At the end of the study (0.5 hr post-administration of VU6021625), plasma and brain samples from each rat (all dose groups) were collected and stored (−80 °C) for bioanalysis by LC-MS/MS as described previously^35,39–40^.

### Abnormal Involuntary Movement Scoring in DRD mice

The dystonia exhibited by knockin mice bearing the c.1160C>A *Th* mutation (DRD mice), which causes DOPA-responsive dystonia in humans, was assessed after treatment with VU6021625. DRD mice were produced as F_1_ hybrids of C57BL/6J +/*Th^DRD^* x DBA/2J +/*Th^DRD^* to avoid the perinatal death associated with inbred C57BL/6J DRD mice^29^. The c.1160C>A mutation in *Th* is coisogenic on C57BL/6 and congenic on DBA/2J. DRD mice were maintained and genotyped as previously described and tested at 16-21 weeks of age^29^. A behavioral inventory was used to define the type of abnormal movement, including tonic flexion or tonic extension (limbs, trunk, head), clonus (limbs), and twisting (trunk, head), as described^38–40^. Homozygous DRD mice (n=8) were habituated to the test cages overnight and experiments started at 8am, when the abnormal movements are most severe in DRD mice^29^. DRD mice were challenged with vehicle or VU6021625 (i.p.) in 1% Tween-80 in a volume of 10 ml/kg. Trihexyphenidyl was used as a positive control. Behavioral assessments began 15 min after compound administration. Abnormal movements were scored for 30 sec at 10 min intervals for 60 min. Mice were tested in a repeated measures design with a pseudorandom order of drug doses and vehicle; each mouse received every dose only once within an experiment. Mice were given at least 4 days between challenges. Observers were blinded to treatment. An arithmetic sum of the disability scores was used to calculate a total score for the entire 1 hr session. These data approximate a continuous variable when total scores from one animal are added together.

## Acknowledgements

Experiments were performed in part through the use of the Neurobehavior Core lab at the Vanderbilt University Medical Center. The authors would like to thank the following funding sources for their generous support of this work: Ancora Innovation, LLC contract to P.J.C. and C.W.L, Michael J. Fox Foundation Target Advancement Program Award 13445 to M.S.M, and NIH grants NIH grants R01NS08852 to E.J.H., R01MH073676 to P.J.C., K99NS110878 to M.S.M. and DoD grant W81XWH-19-1-0355 P.J.C..

## Author Contributions

M.S.M, A.M.B., D.J.F., J.W.D., W.P., T.M.B., Z.K.B., S.C., K.J.W., J.C.O., Y.D. and L.P. performed experiments. M.S.M, A.F.B., D.J.F., J.W.D., W.P., Z.K.B, T.M.B., A.R., C.M.N. analyzed data. M.S.M, D.J.F., D.E., C.M.N., C.W.L, E.J.H, P.J.C., and J.M.R designed experiments. M.S.M and J.M.R wrote the manuscript with the input of all authors.

## Disclosure

C.M.N., J.W.D., C.W.L, P.J.C., T.M.B., A.M.B, J.L.E, and J.M.R. are inventors on applications for composition of matter patents that protect several series of M_4_ antagonists

## Abbreviations

μM: micromolar
5-HT_2B_: serotonin receptor subtype 2B ACh, acetylcholine
ANOVA: analysis of variance
AUC: area under the curve
BG: basal ganglia
cm: centimeter
D_1_: dopamine receptor subtype 1
D_1_-SPN: direct pathway spiny projection neuron
DA: dopamine
DMPK: drug metabolism and pharmacokinetics
DS: dorsal striatum
DRD: DOPA-responsive dystonia
EC_80_: effective concentration of 80% maximal response
ER: efflux ratio
GABA: γ-aminobutyric acid
GPCR: G-protein coupled receptor
HIC: haloperidol induced catalepsy
IC_50_: inhibitory concentration of 50% maximal response
i.p.: intraperitoneal
KO: knockout
L-DOPA: 1-3,4-dihydroxyphenylalanine
M_1_: muscarinic acetylcholine receptor subtype 1
M_2_: muscarinic acetylcholine receptor subtype 2
M_3_: muscarinic acetylcholine receptor subtype 3
M_4_: muscarinic acetylcholine receptor subtype 4
M_5_: muscarinic acetylcholine receptor subtype 5
mAChR: muscarinic acetylcholine receptors
mIPSC: miniature inhibitory post synaptic currents
nAChR: nicotinic acetylcholine receptor
nM: nanomolar
NMR: nuclear magnetic resonance
Oxo-M: oxotremorine
PK: pharmacokinetic
PD: pharmacodynamic
PD: Parkinson Disease
p.o.: per oral
s.c.: subcutaneous
SNc: sunstantia nigra pars compacta
SNr: substantia nigra pars reticulata
WT: wildtype

---

**Scheme 1.**
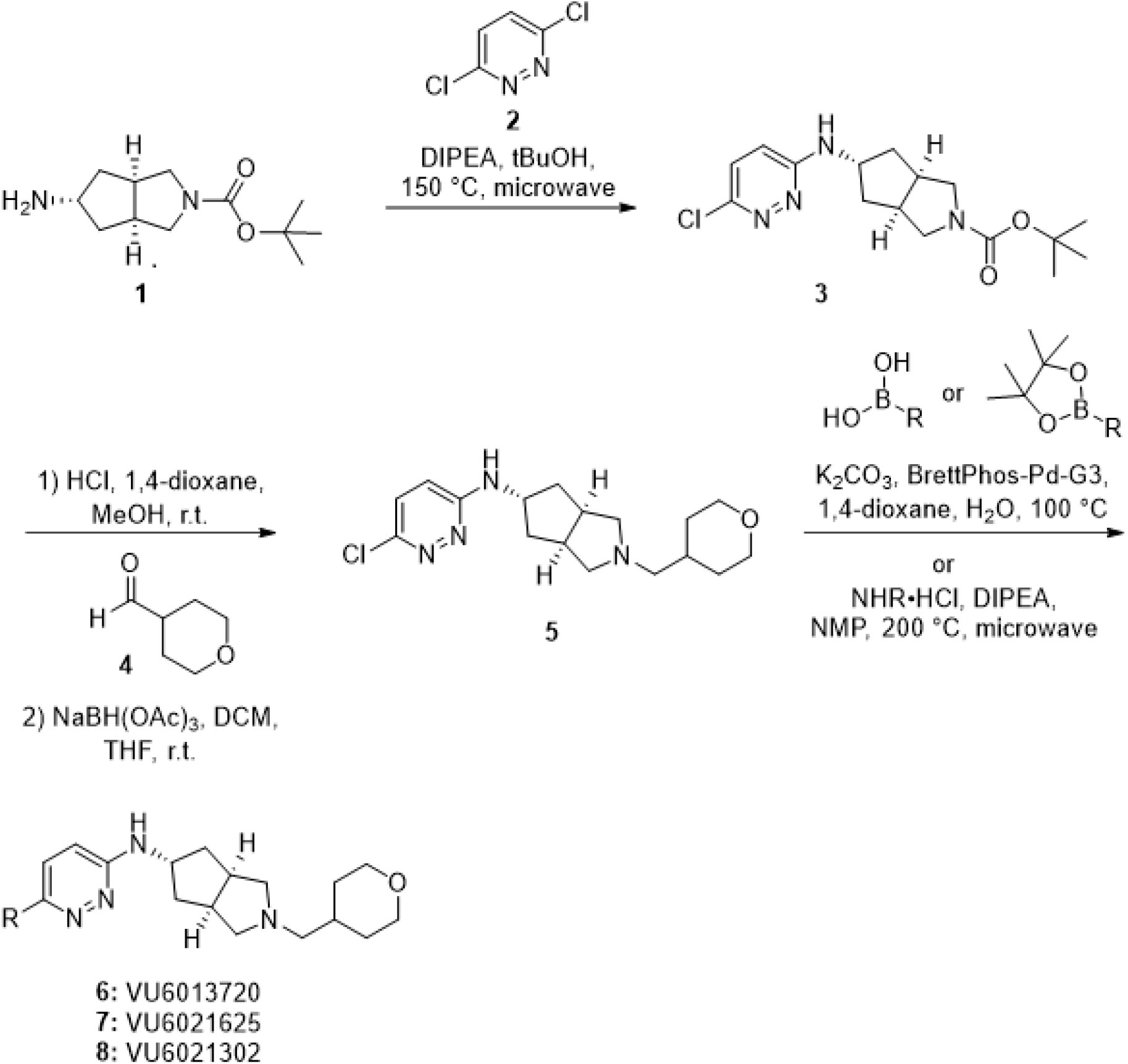

## References

1. Albin RL, Young AB, Penney JB. The functional anatomy of basal ganglia disorders. Trends in neurosciences 1989; 12(10): 366–75.

2. DeLong MR. Primate models of movement disorders of basal ganglia origin. Trends in neurosciences 1990; 13(7): 281–5.

3. DeLong M, Wichmann T. Update on models of basal ganglia function and dysfunction. Parkinsonism & related disorders 2009; 15 Suppl 3(03): S237–40.

4. Nelson AB, Kreitzer AC. Reassessing models of basal ganglia function and dysfunction. Annual review of neuroscience 2014; 37: 117–35.

5. Breakefield XO, Blood AJ, Li Y, Hallett M, Hanson PI, Standaert DG. The pathophysiological basis of dystonias. Nature Reviews Neuroscience 2008; 9(3): 222–34.

6. Karimi M, Perlmutter JS. The role of dopamine and dopaminergic pathways in dystonia: insights from neuroimaging. Tremor and other hyperkinetic movements (New York, NY) 2015; 5: 280.

7. Korczyn AD. Drug treatment of Parkinson’s disease. Dialogues Clin Neurosci 2004; 6(3): 315–22.

8. Thanvi B, Lo N, Robinson T. Levodopa-induced dyskinesia in Parkinson’s disease: clinical features, pathogenesis, prevention and treatment. Postgraduate medical journal 2007; 83(980): 384–8.

9. Moehle MS, Conn PJ. Roles of the M4 acetylcholine receptor in the basal ganglia and the treatment of movement disorders. 2019; 34(8): 1089–99.

10. Conn PJ, Jones CK, Lindsley CW. Subtype-selective allosteric modulators of muscarinic receptors for the treatment of CNS disorders. Trends in pharmacological sciences 2009; 30(3): 148–55.

11. Fahn S, Burke R, Stern Y. Antimuscarinic drugs in the treatment of movement disorders. Progress in brain research 1990; 84: 389–97.

12. Burke RE, Fahn S, Marsden CD. Torsion dystonia: a double-blind, prospective trial of high-dosage trihexyphenidyl. Neurology 1986; 36(2): 160–4.

13. Aosaki T, Miura M, Suzuki T, Nishimura K, Masuda M. Acetylcholine-dopamine balance hypothesis in the striatum: an update. Geriatrics & gerontology international 2010; 10 Suppl 1: S148–57.

14. Benarroch EE. Effects of acetylcholine in the striatum. Recent insights and therapeutic implications. Neurology 2012; 79(3): 274–81.

15. Moehle MS, Pancani T, Byun N, et al. Cholinergic Projections to the Substantia Nigra Pars Reticulata Inhibit Dopamine Modulation of Basal Ganglia through the M(4) Muscarinic Receptor. Neuron 2017; 96(6): 1358–72.e4.

16. Gomeza J, Zhang L, Kostenis E, et al. Enhancement of D1 dopamine receptor-mediated locomotor stimulation in M(4) muscarinic acetylcholine receptor knockout mice. Proc Natl Acad Sci U S A 1999; 96(18): 10483–8.

17. Wess J, Eglen RM, Gautam D. Muscarinic acetylcholine receptors: mutant mice provide new insights for drug development. Nature Reviews Drug Discovery 2007; 6(9): 721–33.

18. Jeon J, Dencker D, Wörtwein G, et al. A subpopulation of neuronal M4 muscarinic acetylcholine receptors plays a critical role in modulating dopamine-dependent behaviors. The Journal of neuroscience. 2010; 30(6): 2396–405.

19. Foster DJ, Wilson JM, Remke DH, et al. Antipsychotic-like Effects of M4 Positive Allosteric Modulators Are Mediated by CB2 Receptor-Dependent Inhibition of Dopamine Release. Neuron 2016; 91(6): 1244–52.

20. Cloud LJ, Jinnah HA. Treatment strategies for dystonia. Expert Opin Pharmacother 2010; 11(1): 5

21. Blokland A, Sambeth A, Prickaerts J, Riedel WJ. Why an M1 Antagonist Could Be a More Selective Model for Memory Impairment than Scopolamine. Frontiers in neurology 2016; 7: 167.

22. Andersson K-E, Campeau L, Olshansky B. Cardiac effects of muscarinic receptor antagonists used for voiding dysfunction. Br J Clin Pharmacol 2011; 72(2): 186–96.

23. Mueller K, Peel JL. Scopolamine produces locomotor stereotypy in an open field but apomorphine does not. Pharmacology, biochemistry, and behavior 1990; 36(3): 613–7.

24. Duty S, Jenner P. Animal models of Parkinson’s disease: a source of novel treatments and clues to the cause of the disease. British journal of pharmacology 2011; 164(4): 1357–91.

25. Croy CH, Chan WY, Castetter AM, Watt ML, Quets AT, Felder CC. Characterization of PCS1055, a novel muscarinic M4 receptor antagonist. European journal of pharmacology 2016; 782: 70–6.

26. Cho TP, Long YF, Gang LZ, et al. Synthesis and biological evaluation of azobicyclo[3.3.0] octane derivatives as dipeptidyl peptidase 4 inhibitors for the treatment of type 2 diabetes. Bioorganic & Medicinal Chemistry Letters 2010; 20(12): 3565–8.

27. Bubser M, Bridges TM, Dencker D, et al. Selective activation of M4 muscarinic acetylcholine receptors reverses MK-801-induced behavioral impairments and enhances associative learning in rodents. ACS Chem Neurosci 2014; 5(10): 920–42.

28. Brady AE, Jones CK, Bridges TM, et al. Centrally active allosteric potentiators of the M4 muscarinic acetylcholine receptor reverse amphetamine-induced hyperlocomotor activity in rats. The Journal of pharmacology and experimental therapeutics 2008; 327(3): 941–53.

29. Rose SJ, Yu XY, Heinzer AK, et al. A new knock-in mouse model of l-DOPA-responsive dystonia. Brain: a journal of neurology 2015; 138(Pt 10): 2987–3002.

30. Kharkwal G, Brami-Cherrier K, Lizardi-Ortiz JE, et al. Parkinsonism Driven by Antipsychotics Originates from Dopaminergic Control of Striatal Cholinergic Interneurons. Neuron 2016; 91(1): 67–78.

31. Shen W, Plotkin JL, Francardo V, et al. M4 Muscarinic Receptor Signaling Ameliorates Striatal Plasticity Deficits in Models of L-DOPA-Induced Dyskinesia. Neuron 2015; 88(4): 762–73.

32. Verbeek DS, Gasser T. Unmet Needs in Dystonia: Genetics and Molecular Biology-How Many Dystonias? Frontiers in neurology 2016; 7: 241.

33. Fuchs T, Saunders-Pullman R, Masuho I, et al. Mutations in GNAL cause primary torsion dystonia. Nature genetics 2013; 45(1): 88–92.

34. Zhao Y, DeCuypere M, LeDoux MS. Abnormal motor function and dopamine neurotransmission in DYT1 DeltaGAG transgenic mice. Experimental neurology 2008; 210(2): 719–30.

35. Downs AM, Fan X, Donsante C, Jinnah HA, Hess EJ. Trihexyphenidyl rescues the deficit in dopamine neurotransmission in a mouse model of DYT1 dystonia. Neurobiology of disease 2019; 125: 115–22.

36. Zhou FM, Lee CR. Intrinsic and integrative properties of substantia nigra pars reticulata neurons. Neuroscience 2011; 198: 69–94.

37. Wenthur CJ, Morrison R, Felts AS, et al. Discovery of (R)-(2-fluoro-4-((-4-methoxyphenyl)ethynyl)phenyl) (3-hydroxypiperidin-1-yl)methanone (ML337), an mGlu3 selective and CNS penetrant negative allosteric modulator (NAM). Journal of medicinal chemistry 2013; 56(12): 5208–12.

38. Devanagondi R, Egami K, LeDoux MS, Hess EJ, Jinnah HA. Neuroanatomical substrates for paroxysmal dyskinesia in lethargic mice. Neurobiology of disease 2007; 27(3): 249–57.

39. Raike RS, Pizoli CE, Weisz C, van den Maagdenberg AMJM, Jinnah HA, Hess EJ. Limited regional cerebellar dysfunction induces focal dystonia in mice. Neurobiology of disease 2013; 49: 200–10.

40. Shirley TL, Rao LM, Hess EJ, Jinnah HA. Paroxysmal dyskinesias in mice. Movement disorders: 2008; 23(2): 259–64.

