## Supplemental Info for "Discovery of the first selective M_4_ muscarinic acetylcholine receptor antagonists with *in vivo* anti-parkinsonian and anti-dystonic efficacy"

**Chemistry Supporting Information**

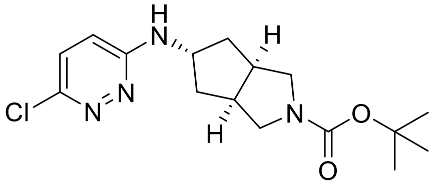


**tert-butyl (3a*R*,5s,6a*S*)-5-((6-chloropyridazin-3-yl)amino)hexahydrocyclopenta[c]pyrrole-2(1H)-carboxylate (3).** Tert-butyl (3a*R*,5s,6a*S*)-5-aminohexahydrocyclopenta[c]pyrrole-2(1H)-carboxylate (5.0 g, 22.1 mmol, 1 eq) and 3,6-dichloropyridazine (9.87 g, 66.3 mmol, 3 eq) were combined in tert-butanol (30 mL), and DIPEA (11.5 mL, 66.3 mmol, 3 eq) was added. The resulting solution was heated to 150 °C under microwave irradiation for 2 h, after which time the reaction mixture was concentrated under reduced pressure, and crude residue was purified by column chromatography (3-100% EtOAc in hexanes) to give the title compound as a white solid (4.87 g, 65%).

$^1$H-NMR (400 MHz, MeOD) δ 7.27 (d, *J* = 9.4 Hz, 1H), 6.87 (d, *J* = 9.4 Hz, 1H), 4.41 (p, *J* = 6.3 Hz, 1H), 3.55 (dd, *J* = 11.1, 8.0 Hz, 2H), 3.19 (dd, *J* = 11.4, 3.8 Hz, 2H), 2.90 – 2.80 (m, 2H), 1.90 – 1.92 (m, 2H), 1.89 – 1.81 (m, 2H), 1.46 (s, 9H).

$^{13}$C-NMR (101 MHz, MeOD) δ 159.3, 156.3, 146.8, 130.3, 120.5, 80.8, 53.7, 53.2 (*signal broadening is observed*) 42.3 (*signal broadening is observed*), 39.5, 28.8.

ES-MS $[M+H]^+$ = 283.2 (- t-butyl).

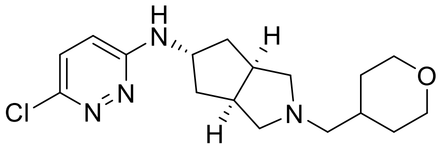


**(3a*R*,5s,6a*S*)-*N*-(6-chloropyridazin-3-yl)-2-((tetrahydro-2H-pyran-4-yl)methyl)octahydrocyclopenta[c]pyrrol-5-amine (5).** Tert-butyl (3a*R*,5s,6a*S*)-5-((6-chloropyridazin-3-yl)amino)hexahydrocyclopenta[c]pyrrole-2(1H)-carboxylate (**3**) (4.86 g, 14.3 mmol, 1 eq) was dissolved in 1,4-dioxane (70 mL) and MeOH (20 mL), and 4M HCl in dioxanes solution (50 mL) was added dropwise. The resulting solution was stirred at r.t. for 1 h, after which time solvents were concentrated under reduced pressure to give the HCl salt as a white solid, which was dried under vacuum and used without additional purification (3.95 g, 100%). The HCl salt was then suspended in DCM (40 mL) and THF (50 mL), and tetrahydro-2H-pyran-4-carbaldehyde (2.29 g, 20.1 mmol, 1.4 eq) was added, followed by sodium triacetoxyborohydride (6.08 g, 28.9

mmol, 2 eq). The resulting solution was stirred at r.t. for 1.5 h, after which time the reaction mixture was quenched with sat. $NaHCO_3$, and extracted with DCM. Combined organic extracts were washed with brine, and dried over $MgSO_4$. Solvents were filtered and concentrated to give the title compound as a white solid (4.31 g, 89% over 2 steps).

$^1$H-NMR (400 MHz, MeOD) δ 7.26 (d, *J* = 9.4 Hz, 1H), 6.86 (d, *J* = 9.4 Hz, 1H), 4.43 – 4.36 (m, 1H), 3.93 (dd, *J* = 11.3, 3.7 Hz, 2H), 3.42 (td, *J* = 11.9, 1.9 Hz, 2H), 2.84 – 2.68 (m, 4H), 2.31 (d, *J* = 6.8 Hz, 2H), 2.27 – 2.17 (m, 2H), 1.91 (ddd, *J* = 12.9, 5.9, 2.1 Hz, 2H), 1.83 – 1.64 (m, 5H), 1.32 – 1.21 (m, 2H).

$^{13}$C-NMR (101 MHz, MeOD) δ 159.7, 146.6, 130.3, 120.4, 68.9, 63.5, 62.9, 53.4, 41.5, 39.1, 35.3, 32.9.

ES-MS $[M+H]^+$ = 337.2.

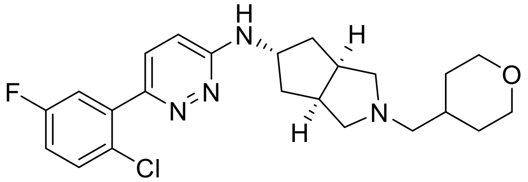


**(3a*R*,5s,6a*S*)-*N*-(6-(2-chloro-5-fluorophenyl)pyridazin-3-yl)-2-((tetrahydro-2H-pyran-4-yl)methyl)octahydrocyclopenta[c]pyrrol-5-amine (6, VU6013720)** (3a*R*,5s,6a*S*)-*N*-(6-chloropyridazin-3-yl)-2-((tetrahydro-2H-pyran-4-yl)methyl)octahydrocyclopenta[c]pyrrol-5-amine (**5**) (100 mg, 0.30 mmol, 1 eq), 2-chloro-5-fluorophenylboronic acid (62 mg, 0.36 mmol, 1.2 eq), potassium carbonate (125 mg, 0.89 mmol, 3 eq) and BrettPhos-Pd-G3 (27 mg, 0.030 mmol, 0.1 eq) were combined in a vial, and 5:1 1,4-dioxane/$H_2O$ solution (5 mL total, degassed under vacuum) was added via syringe. The resulting mixture was stirred under an inert atmosphere at 100 °C for 2.5 h, after which time the reaction mixture was cooled to r.t. and diluted with water and DCM. The aqueous layer was extracted with DCM, and combined organic extracts were filtered through a phase separator and concentrated. Crude residue was purified by RP-HPLC (10-50% MeCN in 0.1% TFA aqueous solution over 20 min). Fractions containing product were basified with sat. $NaHCO_3$, and extracted with DCM. Combined organic extracts were dried over $MgSO_4$, filtered and concentrated under reduced pressure to give the title compound as a white solid (40 mg, 31%).

$^1$H-NMR (400 MHz, MeOD) δ 7.57 – 7.52 (m, 2H), 7.35 (dd, *J* = 9.0, 3.1 Hz, 1H), 7.20 (ddd, *J* = 8.8, 7.9, 3.1 Hz, 1H), 6.91 (d, *J* = 9.3 Hz, 1H), 4.56 – 4.48 (m, 1H), 3.94 (dd, *J* = 11.2, 3.5 Hz, 2H), 3.43 (td, *J* = 11.9, 1.9 Hz, 2H), 2.87 – 2.74 (m, 4H), 2.34 (d, *J* = 6.9 Hz, 2H), 2.26 (dd, *J* = 8.4, 4.0 Hz, 2H), 1.97 (ddd, *J* = 12.9, 5.9, 2.1 Hz, 2H), 1.84 – 1.70 (m, 5H), 1.34 – 1.22 (m, 2H).

$^{13}$C-NMR (101 MHz, MeOD) δ 162.8 (d, $J$ = 246.1 Hz), 159.7, 150.95 (d, $J$ = 1.9 Hz), 139.8 (d, $J$ = 8.1 Hz), 132.8 (d, $J$ = 8.5 Hz), 130.5, 128.6 (d, $J$ = 3.3 Hz), 118.9 (d, $J$ = 24.1 Hz), 118.0 (d, $J$ = 23.0 Hz), 116.2, 68.9, 63.6, 63.0, 53.3, 41.5, 39.3, 35.3, 33.0.

ES-MS $[M+H]^+$ = 431.4.

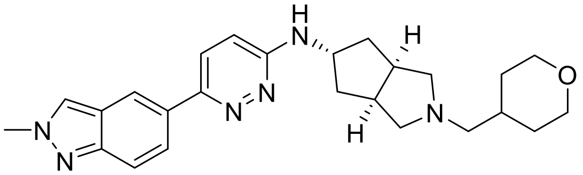

**(3a*R*,5s,6a*S*)-*N*-(6-(2-methyl-2H-indazol-5-yl)pyridazin-3-yl)-2-((tetrahydro-2H-pyran-4-yl)methyl)octahydrocyclopenta[c]pyrrol-5-amine (7, VU6021625).** (3a*R*,5s,6a*S*)-*N*-(6-chloropyridazin-3-yl)-2-((tetrahydro-2H-pyran-4-yl)methyl)octahydrocyclopenta[c]pyrrol-5-amine (**5**) (1.0 g, 2.97 mmol, 1 eq), 2-methylindazole-5-boronic acid pinacol ester (996 mg, 3.86 mmol, 1.3 eq), potassium carbonate (1.25 g, 8.91 mmol, 3 eq) and BrettPhos-Pd-G3 (269 mg, 0.30 mmol, 0.1 eq) were combined in a vial, and 5:1 1,4-dioxane/$H_2O$ solution (15 mL total, degassed under vacuum) was added via syringe. The resulting mixture was stirred under an inert atmosphere at 100 °C for 3 h, after which time the reaction mixture was cooled to r.t. and diluted with water and DCM. The aqueous layer was extracted with DCM, and combined organic extracts were dried over $MgSO_4$, filtered and concentrated under reduced pressure. Crude residue was purified by RP-HPLC (20-60% MeCN in 0.05% $NH_4OH$ aqueous solution over 20 min). Fractions containing product were concentrated to give the title compound as a white solid (431 mg, 34%).

$^{1}$H-NMR (400 MHz, MeOD) δ 8.27 (s, 1H), 8.17 (dd, $J$ = 1.7, 0.9 Hz, 1H), 7.95 (dd, $J$ = 9.1, 1.7 Hz, 1H), 7.78 (d, $J$ = 9.4 Hz, 1H), 7.67 (dt, $J$ = 9.1, 1.0 Hz, 1H), 6.93 (d, $J$ = 9.4 Hz, 1H), 4.54 – 4.47 (m, 1H), 4.23 (s, 3H), 3.94 (dd, $J$ = 11.0, 3.4 Hz, 2H), 3.43 (td, $J$ = 11.9, 2.0 Hz, 2H), 2.91 – 2.73 (m, 4H), 2.36 (d, $J$ = 6.9 Hz, 2H), 2.30 – 2.26 (m, 2H), 2.00 – 1.95 (m, 2H), 1.85 – 1.70 (m, 5H), 1.33 – 1.22 (m, 2H).

$^{13}$C-NMR (101 MHz, MeOD) δ 159.3, 152.5, 150.1, 131.9, 127.4, 127.3, 126.3, 123.6, 119.0, 117.8, 68.9, 63.5, 62.9, 53.3, 41.5, 40.3, 39.2, 35.2, 32.9. *Note: 1 aromatic signal is obscured.*

ES-MS $[M+H]^+$ = 433.0.

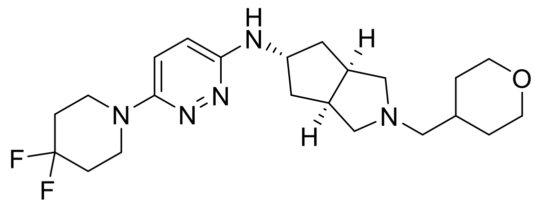

**(3a*R*,5s,6a*S*)-*N*-(6-(4,4-difluoropiperidin-1-yl)pyridazin-3-yl)-2-((tetrahydro-2H-pyran-4-yl)methyl)octahydrocyclopenta[c]pyrrol-5-amine (8, VU6021302).** (3a*R*,5s,6a*S*)-*N*-(6-chloropyridazin-3-yl)-2-((tetrahydro-2H-pyran-4-yl)methyl)octahydrocyclopenta[c]pyrrol-5-amine (**5**) (1.0 g, 2.96 mmol, 1 eq) and 4,4-difluoropiperidine hydrochloride (4.68 g, 29.7 mmol, 10 eq) were combined in NMP (10 mL), and DIPEA (5.17 mL, 29.7 mmol, 10 eq) was added. The resulting solution was stirred under microwave irradiation at 200 °C for 2 h, after which time the reaction mixture was purified directly by RP-HPLC (25-65% MeCN in 0.05% $NH_4OH$ aqueous solution over 20 min). Fractions containing product were concentrated to give the title compound as a slightly tan solid (797 mg, 64%).

$^1$H-NMR (400 MHz, MeOD) δ 7.18 (d, *J* = 9.7 Hz, 1H), 6.81 (d, *J* = 9.7 Hz, 1H), 4.36 – 4.29 (m, 1H), 3.93 (dd, *J* = 11.1, 3.4 Hz, 2H), 3.57 – 3.54 (m, 4H), 3.42 (td, *J* = 11.8, 2.0 Hz, 2H), 2.88 – 2.86 (m, 2H), 2.79 – 2.69 (m, 2H), 2.34 (d, *J* = 6.9 Hz, 2H), 2.23 (dd, *J* = 9.3, 5.1 Hz, 2H), 2.08 – 1.98 (m, 4H), 1.91 (ddd, *J* = 12.9, 6.1, 2.3 Hz, 2H), 1.83 – 1.62 (m, 5H), 1.31 – 1.21 (m, 2H).

$^{13}$C-NMR (101 MHz, MeOD) δ 155.9, 155.7, 123.3 (t, *J* = 240.8 Hz), 120.8, 120.7, 68.9, 63.5, 62.9, 53.4, 45.2 (t, *J* = 5.2 Hz), 41.4, 39.2, 35.2, 34.3 (t, *J* = 23.0 Hz), 32.9.

ES-MS $[M+H]^+$ = 422.5.

Supplementary Table 1

| Target | Radioligand | Species | % Inhibition |
|---|---|---|---|
| Adenosine A1 | [3H] DPCPX | Human | 12 |
| Adenosine A2A | [3H] CGS-21680 | Human | 5 |
| Adenosine A3 | [125I] AB-MECA | Human | 10 |
| Adrenergic a1A | [3H] Prazosin | Rat | 13 |
| Adrenergic a1B | [3H] Prazosin | Rat | 3 |
| Adrenergic a1D | [3H] Prazosin | Human | 17 |
| Adrenergic a2A | [3H] Rauwolscine | Human | 18 |
| Adrenergic ß1 | [125I] Cyanopindolol | Human | 6 |
| Adrenergic ß2 | [3H] CGP-12177 | Human | 0 |
| Androgen (Testosterone) | [3H] Methyltrienolone | Human | -13 |
| Bradykinin B1 | [3H] (Des-Arg10, Leu9)- Kallidin | Human | -10 |
| Bradykinin B2 | [3H] Bradykinin | Human | -7 |
| Calcium Channel L-Type, Benzothiazepine | [3H] Diltiazem | Human | 7 |
| Calcium Channel L-Type, Dihydropyridine | [3H] Nitrendipine | Rat | 1 |
| Calcium Channel N-Type | [125I] ω-Conotoxin GVIA | Rat | -9 |
| Cannabinoid CB1 | [3H] SR141716A | Rat | -3 |

| | | | |
|---|---|---|---|
| Dopamine D1 | [3H] SCH-23390 | Human | 2 |
| Dopamine D2S | [3H] Spiperone | Human | 17 |
| Dopamine D3 | [3H] Spiperone | Human | 34 |
| Dopamine D4.4 | [3H] Spiperone | Human | 3 |
| Endothelin ETA | [125I] Endothelin-1 | Human | -11 |
| Endothelin ETB | [125I] Endothelin-1 | Human | 2 |
| Epidermal Growth Factor (EGF) | [125I] EGF | Human | -5 |
| Estrogen ERa | [3H] Estradiol | Human | -3 |
| GABAA, Flunitrazepam, Central | [3H] Flunitrazepam | Rat | -4 |
| GABAA, Muscimol, Central | [3H] Muscimol | Rat | -11 |
| GABAB1A | [3H] CGP-54626 | Human | 2 |
| Glucocorticoid | [3H] Dexamethasone | Human | -2 |
| Glutamate, Kainate | [3H] Kainic acid | Rat | 6 |
| Glutamate, NMDA, Agonism | [3H] CGP-39653 | Rat | -5 |
| Glutamate, NMDA, Glycine | [3H] MDL 105,519 | Rat | -12 |
| Glutamate, NMDA, Phencyclidine | [3H] TCP | Rat | -1 |
| Histamine H1 | [3H] Pyrilamine | Human | 25 |
| Histamine H2 | [125I] Aminopotentidine | Human | -3 |
| Histamine H3 | [3H] N-α-Methylhistamine | Human | 88 |
| Imidazoline I2, Central | [3H] Idazoxan | Rat | 2 |

| | | | |
|---|---|---|---|
| Interleukin IL-1 R1 | [125I] Interleukin-1β | Human | -7 |
| Leukotriene, Cysteinyl CysLT1 | [3H] LTD4 | Human | 12 |
| Melatonin MT1 | [125I] 2-Iodomelatonin | Human | -2 |
| Muscarinic M1 | [3H] N-Methylscopolamine | Human | 33 |
| Muscarinic M2 | [3H] N-Methylscopolamine | Human | 85 |
| Muscarinic M3 | [3H] N-Methylscopolamine | Human | 51 |
| Neuropeptide Y Y1 | [125I] Peptide YY | Human | -13 |
| Neuropeptide Y Y2 | [125I] Peptide YY | Human | -2 |
| Nicotinic Acetylcholine α1, Bungarotoxin | [125I] α-Bungarotoxin | Human | -1 |
| Nicotinic Acetylcholine α3β4 | [125I] Epibatidine | Human | 55 |
| Opiate δ1 (OP1, DOP) | [3H] Naltrindole | Human | -1 |
| Opiate κ (OP2, KOP) | [3H] Diprenorphine | Human | 5 |
| Opiate μ (OP3, MOP) | [3H] Diprenorphine | Human | 6 |
| Phorbol Ester | [3H] PDBu | Mouse | 9 |
| Platelet Activating Factor (PAF) | [3H] PAF | Human | -4 |
| Potassium Channel [KATP] | [3H] Glyburide | Human | 11 |
| Potassium Channel hERG | [3H] Astemizole | Human | 23 |

| | | | |
|---|---|---|---|
| Prostanoid EP4 | [3H] Prostaglandin E2 | Human | 0 |
| Purinergic P2X | [3H] α, β-Methylene-ATP | Rat | 4 |
| Rolipram | [3H] Rolipram | Rat | 0 |
| Serotonin (5-Hydroxytryptamine) 5-HT1A | [3H] 8-OH-DPAT | Human | 6 |
| Serotonin (5-Hydroxytryptamine) 5-HT2B | [3H] Lysergic acid diethylamide | Human | 53 |
| Serotonin (5-Hydroxytryptamine) 5-HT3 | [3H] GR-65630 | Human | -2 |
| Sigma σ1 | [3H] Haloperidol | Human | 38 |
| Sodium Channel, Site 2 | [3H] Batrachotoxinin | Rat | 15 |
| Tachykinin NK1 | [3H] Substance P | Human | -10 |
| Thyroid Hormone | [125I] Triiodothyronine | Rat | -21 |
| Transporter, Dopamine (DAT) | [125I] RTI-55 | Human | 15 |
| Transporter, GABA | [3H] GABA | Rat | -5 |
| Transporter, Norepinephrine (NET) | [125I] RTI-55 | Human | 15 |
| Transporter, Serotonin (5-Hydroxytryptamine) (SERT) | [3H] Paroxetine | Human | -1 |
